# Mosquito thermal tolerance is remarkably constrained across a large climatic range

**DOI:** 10.1101/2023.03.02.530886

**Authors:** Lisa I. Couper, Johannah E. Farner, Kelsey P. Lyberger, Alexandra S. Lee, Erin A. Mordecai

## Abstract

How mosquitoes may respond to rapid climate warming remains unknown for most species, but will have major consequences for their future distributions, with cascading impacts on human well-being, biodiversity, and ecosystem function. We investigated the adaptive potential of a wide-ranging mosquito species, *Aedes sierrensis*, across a large climatic gradient by conducting a common garden experiment measuring the thermal limits of mosquito life history traits. Although field-collected populations originated from vastly different thermal environments that spanned over 1,200 km, we found remarkably limited variation in upper thermal tolerance between populations, with the upper thermal limits of fitness varying by <1°C across the species range. For one life history trait—pupal development rate—we did detect significant variation in upper thermal limits between populations, and this variation was strongly correlated with source temperatures, providing evidence of local thermal adaptation for pupal development. However, we found environmental temperatures already regularly exceed our highest estimated upper thermal limits throughout most of the species range, suggesting limited potential for mosquito thermal tolerance to evolve on pace with warming. Strategies for avoiding high temperatures such as diapause, phenological shifts, and behavioral thermoregulation are likely important for mosquito persistence.

## Introduction

How mosquitoes respond in the face of rapid anthropogenic climate warming is a key open question of ecological and public health concern. As temperature impacts nearly all aspects of mosquito life cycles, climate warming may cause large shifts in their distributions and dynamics^1,2^. In particular, current predictions suggest that mosquito distributions may shift higher in latitude and elevation, expanding into temperate regions as they become newly suitable, and contracting in some tropical regions as they become too warm^3–6^. However, these predictions have not typically incorporated the potential for mosquito adaptive responses, and thus may overestimate declines at current warm edges.

Temperature sets fundamental limits on mosquito distributions as mosquito survival and reproduction are inhibited beyond critical thermal limits. As temperatures exceed those limits under warming, mosquito populations could persist through a variety of mechanisms including range shifts to track suitable temperatures, shifts in daily and/or seasonal activity patterns to avoid high temperatures, behavioral thermoregulation (*i.e.,* actively seeking out cooler microhabitats), and increased heat tolerance through evolutionary adaptation^7^. Of these responses, evolutionary adaptation may be particularly important for enabling long-term persistence, but the potential for mosquito thermal adaptation remains poorly understood, owing to several empirical knowledge gaps^8–10^.

A key component of whether a given mosquito species can evolutionarily adapt to warming is the presence of standing variation in upper thermal tolerance within a species^10^. Decades of research on mosquito thermal biology have demonstrated variation in thermal performance between species (*e.g.*^1,11,12^). Further, several studies have identified within-species variation in response to other aspects of climate, such as cold tolerance in *Aedes albopictus*^13,14^ and aridity tolerance in *Anopheles gambiae*^15,16^. Only a few studies have investigated within-species variation in upper thermal tolerance, and have generally found some evidence of standing variation (*i.e*., differing rates of survival, reproduction, or development among populations at high temperatures), but little evidence of local *thermal adaptation* (*i.e.*, higher heat tolerance observed in populations from warmer environments than those from cooler environments)^17–21^. However, these studies typically investigated relatively few mosquito populations from a limited portion of the species range, owing to logistical challenges of collecting, rearing, and experimenting on many wide-ranging populations. Further, mosquito thermal tolerance was typically measured on select life history traits or metabolic rates, potentially obscuring patterns of thermal adaptation evident across the full life cycle^9,22,23^. Thus, the extent of variation in upper thermal tolerance among populations within a species and the evidence for thermal adaptation is still unknown.

We set out to rigorously investigate the evidence for mosquito thermal adaptation by using *Aedes sierrensis*, the western tree hole mosquito, as a novel model system. *Ae. sierrensis* makes an ideal model species for this investigation because it is commonly occurring across its distribution (ranging from Southern California to British Columbia and coastal to montane environments^24,25^), which covers a large range of thermal environments, presenting varying selection pressures and opportunities for local thermal adaptation. This species has a seasonal life cycle driven by temperature, precipitation, and day length cues, and which occurs in discrete, easy-to-sample habitat (water-filled tree holes)^25^, facilitating field collection of individuals at the same life stage across the species range. Further, although *Ae. sierrensis* is not a known vector of human pathogens, it is congeneric to major human disease vectors (*i.e., Ae. aegypti, Ae. albopictus*) and is itself a vector of dog heartworm, making results potentially informative for understanding warming responses in these vector species. Leveraging this model system, we set out to answer the following specific research questions: (i) How much does thermal tolerance vary between populations across the species range? (ii) Is variation in thermal tolerance, if observed, correlated with the source thermal environment? (*i.e.,* is there evidence of local thermal adaptation?) (iii) At present, how often do environmental temperatures exceed mosquito populations’ upper thermal limits?

To answer these questions, we conducted a common garden experiment using ten *Ae. sierrensis* populations spanning nearly the entire species range (1,200 km; Figure 1). The thermal environments of collected populations varied widely, with annual mean temperatures varying by >7°C, and average daily maxima in the spring and summer varying by >5°C. We reared these field-collected populations in the lab for one generation at common temperatures, then separated F1 individuals into one of six temperature treatments ranging from 5-32LJ. We tracked individuals daily to approximate individual-level fitness, as well as its component life history traits—larval and pupal survival and development rates, adult lifespan, and wing length (a proxy for fecundity). We then fit thermal performance curves to these experimental data to estimate upper and lower thermal limits, thermal optima, and breadth, and maximum performance for each population and trait. In our investigation of variation in mosquito thermal tolerance, we compared variation in these estimated upper thermal limits for each trait and population. We note that prior studies of mosquito thermal tolerance have used a variety of methods to measure thermal tolerance including static and dynamic heat tolerance assays (*e.g*., ‘thermal knockdowns’)^12,26^, reciprocal transplants^13^, and comparisons of niche-based distribution models^27^. These methods may each capture a slightly different component of thermal tolerance (*e.g.*, capacity for heat shock responses, combined genetic and plastic responses), thus our metric of thermal tolerance may not be comparable across all approaches. We focused on upper thermal limits from trait thermal performance curves as they capture high temperature constraints across the life span.

**Figure 1.**
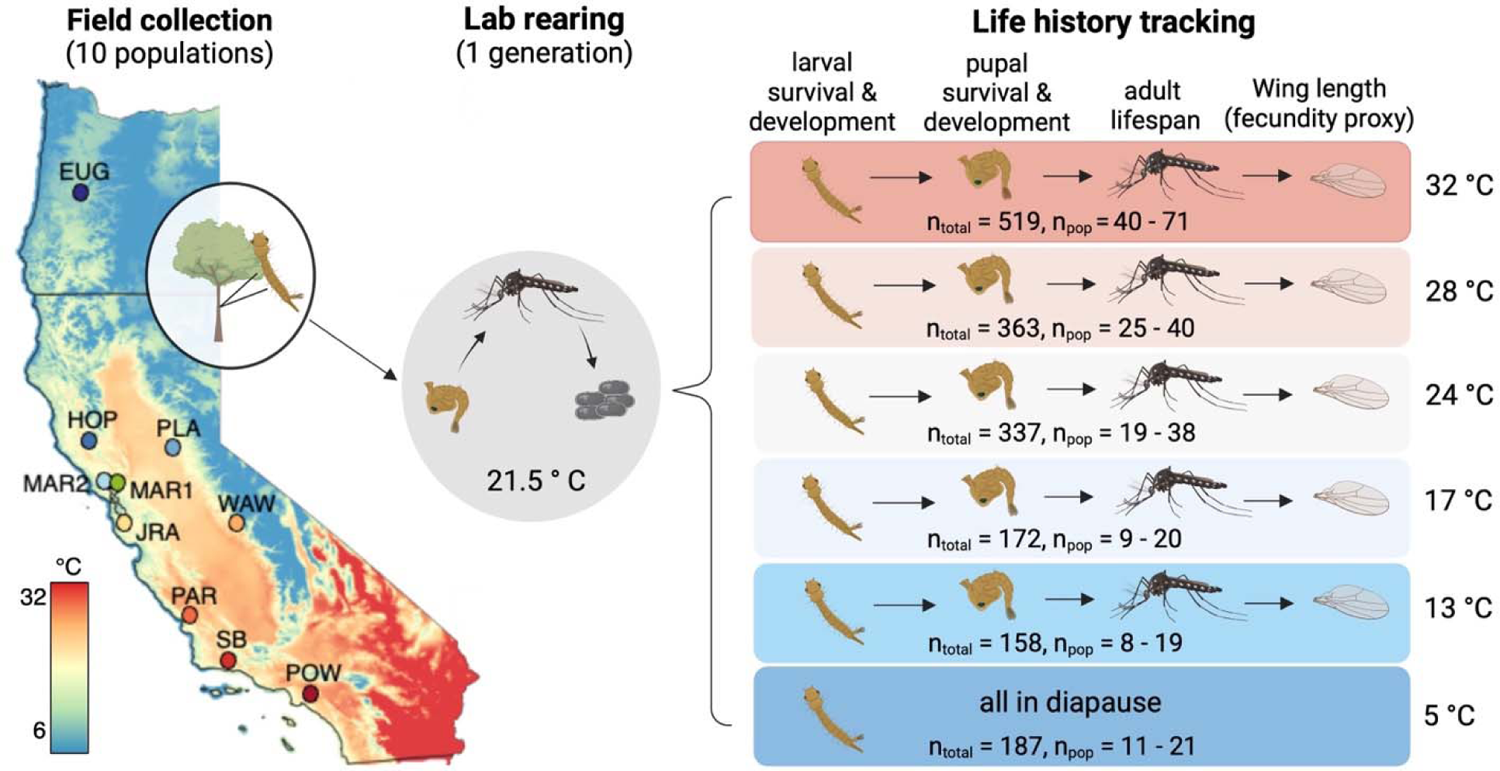
Sample collection locations and experimental design used to measure mosquito thermal performance. Ten populations were collected as larvae from tree holes across the Western U.S. and reared in the lab under common conditions for one generation. The resulting larvae from each population were randomly designated into one of six temperature treatments. The total number of larvae assigned to each treatment is noted above (‘n_total_’) as is the range of larvae from each population (‘n_pop_’); Supplemental Table S2 indicates the full breakdown of larvae per population and treatment). Individuals were checked daily for life stage transitions (*e.g.*, larvae to pupae, pupae to adult) or death. Map colors denote the average maximum annual temperature (LJ) from 1991 – 2020 from PRISM data. Supplemental Figure S1 shows the average minimum and mean temperature across this same extent. Population metadata, including full site names, latitude, longitude, and elevation are provided in Supplemental Table S1.

Despite originating from a wide range of thermal environments, populations differed very little in their thermal limits for fitness, and for nearly all other life history traits. For pupal development rate, we did find significant variation in upper thermal limits between populations, with five times greater variation in upper thermal limits than previously found in ectotherm species across this same range. Further, this variation corresponded with populations’ source thermal environments, providing evidence of local thermal adaptation. However, environmental temperatures across most of the species range already regularly exceed populations’ estimated upper thermal limits, suggesting thermal adaptation alone may play a limited role in enabling persistence under warming. Seasonal life history strategies and behavioral thermoregulation are likely important strategies for mosquitoes coping with ongoing climate warming.

## Methods

### Field collection

*Ae. sierrensis* typically completes one life cycle per year, with adults laying eggs in naturally occurring tree holes. Eggs hatch when the tree holes fill with water beginning in the late fall and advance through four larval instars and one pupal life stage throughout the winter before eclosing as adults in the spring and summer^24^. Most North American *Ae. sierrensis* populations (*e.g.*, those from 26-46°N), including all of our collected populations, undergo diapause between the fourth larval instar and pupal life stage, and all populations undergo embryonic diapause^28^. We collected larval *Ae. sierrensis* from 346 tree holes spanning over 1,200 km across the Western U.S. between October 2021 and April 2022 (Figure 1, Supplemental Table S1 for collection metadata). We collected *Ae. sierrensis* and tree hole water in plastic cups and maintained these at cold temperatures (< 10°C) during transportation to the lab, then at 4LJ until processing. We visually inspected individuals from each sampled tree hole for the presence of *Lambornella clarki*—a ciliate parasite that can infect larval *Ae. sierrensis*. Only larvae from tree holes without the parasite were used in this experiment. Further, to maintain sufficient genetic variation and avoid excessive inbreeding, we reared only larvae from tree holes with at least 30 collected individuals.

### Lab rearing

After processing, we maintained select populations (*i.e.*, those from tree holes with ≥30 individuals and no *L. clarki*) under shared lab conditions of 21.5LJ, and a 13 h: 11 h light:dark cycle. We periodically fed larvae a finely-ground mix of Tetramin fish flakes (48% by weight), guinea pig chow (48%), and liver powder (2%). Once reaching the adult stage, we housed populations in 8 x 8 x 8 cm aluminum collapsible cages (BioQuip, Rancho Dominguez, CA, USA) with continuous access to a 10% sugar solution. We offered each population a blood meal of defibrinated sheep’s blood approximately once per week and placed an oviposition cup, consisting of a paper cup lined with water-soaked coffee filter paper, inside each cage within four days of the first blood-feeding. We collected eggs and held these at room temperature for two weeks, then in the refrigerator at 4LJ and near 24 h darkness to mimic winter conditions and promote hatching (potentially because these cold, dark conditions cause eggs to enter and exit diapause, as would occur in the field; pers. comm. Bret Barner, Solano County vector control), which occurred 1-3 months later.

To ensure sufficient sample sizes for each treatment of the experiment, we only used populations that produced >300 eggs in total. This resulted in 10 populations for use in the experiment (Figure 1), wherein ‘population’ refers to a group of individuals originating from the same tree hole. These collections are highly likely to represent distinct populations, as the minimum distance between any pair of populations used in the experiment was 3.4 km, and *Ae. sierrensis* adults are weak fliers and typically do not disperse far from their larval tree hole^29^. We note that a more precise definition of a population would incorporate specific dispersal capabilities and/or genetic structuring, but this has not yet been investigated for *Ae. sierrensis*.

To hatch eggs, we prepared a separate tray for each population, which consisted of 500 mL Arrowhead distilled water, 300 mL autoclaved tree hole water (combined from all sampled tree holes), and ¾ tsp Brewers’ yeast. We submerged egg papers from each population in trays between July 4 - 6, 2022, 24 h after the respective hatching tray was prepared.

We note that by using F1 individuals in our experiment, we have not eliminated maternal/cross-generational effects, which may impact thermal tolerance^30^. That is, while we sought to minimize direct environmental effects on thermal tolerance (*i.e*., ‘phenotypic plasticity’) and capture genetically-based differences, environmental effects from prior generations could still impact F1 thermal tolerance.

### Experimental design

The experiment consisted of tracking life histories for individual *Ae. sierrensis* from one of ten populations, held at one of six temperature treatments (Figure 1; see Supplemental Table S2 for sample sizes). The temperature treatments–5, 13, 17, 24, 28, and 32LJ–were chosen based on the range of temperatures realistically experienced by *Ae. sierrensis* in the field and based on survival rates assessed during pilot experiments conducted in the lab (Figure 1, Supplemental Figure S1). These constant temperatures were maintained using Fisher Scientific Isotemp incubators (for the 13, 24, 28, and 32LJ treatments) and climate-controlled rooms (for the 5 and 17LJ treatments). Although fluctuating temperatures could have more closely mimicked natural conditions, we chose to use constant temperatures here as it provides a baseline for characterizing thermal responses and because measuring all possible combinations of temperature mean and variability would have been intractable.

The experiment began with larvae emerging 48 h after egg paper submersion (*i.e.*, approximately 1-day old larvae). For each individual, we measured the following traits: larval survival, larval development rate, pupal survival, pupal development rate, adult lifespan, and wing length (a proxy for fecundity; see *Methods: Measuring wing length as a proxy for fecundity*). We intentionally included more larvae from each population in the higher temperature treatments as we expected greater mortality at these temperatures based on pilot experiments. We visually inspected each individual on a daily basis, recording life stage transitions and deaths, and moving individuals into the appropriate housing for the given life stage. We maintained larvae in plastic containers in groups of five with approximately 100 mL of water and 4 mg of larval food, in accordance with *Aedes* rearing protocols that promote high larval survivorship in the absence of other factors^31,32^. We maintained pupae individually in glass vials with approximately 5 mL deionized water. Upon eclosion, we transferred adults to individual 4 oz plastic specimen cups with one 10% sugar-soaked cotton ball and observed each individual until death. To estimate fecundity of individuals that died as adults, we removed and measured the length of their left wing, a commonly used proxy^33,34^ (see *Methods: Measuring wing length as a proxy for fecundity*). Any larva that was alive but had not pupated by September 28, 2022 (*i.e.*, 82-84 days after larval emergence) was counted as survived for the larval survival trait and considered to be in diapause.

### Measuring wing length as a proxy for fecundity

To estimate individual fecundity, we measured the wing length of each individual used in the experiment. While wing length is an imperfect measure of fecundity, it is widely used in the literature and has been validated for several mosquito species (*e.g.*, *Anopheles arabiensis*^33^, *Anopheles gambiae*^34^, *Aedes albopictus*^35–37^, *Aedes geniculatus*^38^) in addition to *Aedes sierrensis*^39^. Further, using this proxy enabled us to obtain both a lifespan and estimate of reproductive output for each individual used in the experiment, whereas individually blood-feeding hundreds of mosquitoes held inside incubators would have been intractable. To measure wing length, we removed and photographed the left wing mounted on a microscope slide with a 1 mm graticule. We then used ImageJ to measure the wing length as the distance from the alular notch to the tip of the wing margin excluding the fringe scales, using the 1 mm graticule for calibration^40^ (Supplemental Figure S2). We then used the relationship between *Ae. sierrensis* female wing length and the number of eggs laid in the first clutch established in Washburn et al. 1989^39^ (*i.e.*, 51.33 x female wing length (mm) - 87.96). We validated this relationship in the lab using a separate, smaller number of individuals from our experimental populations (see Supplemental Methods; Supplemental Figure S3).

### Fitness estimation

We estimated an individual-level mosquito fitness proxy—here defined as a measure of individual reproductive output through the first gonotrophic cycle—as survival to reproductive maturity multiplied by estimated fecundity in the first gonotrophic cycle. For survival to reproductive maturity, we considered whether an individual survived to adulthood and achieved an adult lifespan of 10 days at 24 or 28LJ, 11 days at 17LJ, or 17 days at 13LJ. These lifespans represent the minimum number of days from adult eclosion to egg-laying at a given temperature, as observed in the validation experiment (*Supplemental Methods: Determining age at reproductive maturity*). As no individuals eclosed at 5LJ and no individuals survived longer than one day at 32LJ, all individuals at these two temperature treatments were estimated to have zero fitness. For estimated fecundity, we used the wing length approximation described above (*Methods: Measuring wing length as a proxy for fecundity*). As these estimates were made for both males and females, we multiplied the estimated fecundity of a given adult by the proportion of females from that population and temperature treatment.

### Characterizing the source thermal environment

We characterized the source thermal environment of each population using climate data from PRISM, which we accessed and analyzed using Google Earth Engine^41^. PRISM provides gridded climate data at a 4 km resolution by downscaling data from a network of monitoring stations^42^. We used either daily or monthly temperature data from 2000 – 2020 to calculate key variables capturing temperature means, variations, and extremes. We specifically sought to include only biologically meaningful temperature variables, such as those previously associated with thermal tolerance in ectotherms^43^, rather than many possible characterizations of climate (*e.g.,* all 19 WorldClim bioclimatic variables). These variables included annual mean temperature, mean temperature in January – March (the period when eggs typically exit diapause and hatch as larvae), seasonal variation in temperature (defined as the difference between the mean warmest month temperature and the mean coolest month temperature), average warm-season maximum (defined as the mean daily maxima in the Spring and Summer), and the number of days where maximum temperatures exceeded 35LJ (the highest upper thermal limit for any trait estimated from our experimental data) excluding periods of potential dormancy (*e.g.,* August – October).

Variables were calculated at a 1 km buffer around the sampled tree hole for each population, approximating the geographic range of an individual mosquito. We investigated Pearson’s correlations between these temperature variables and select thermal performance parameters and traits (*i.e.,* those with significant between-population variation).

While the above estimates of source environmental temperature likely capture the thermal conditions for populations at a broad spatial scale, they may not reflect the exact temperatures within a given tree hole. We sought to directly measure tree hole temperatures for each population by placing iButton temperature loggers (DS1921G, manufactured by Maxim Integrated, San Jose, California) in each sampled tree hole at the time of location; however, only two iButtons were recovered the following year. For these two tree holes, we compare the direct temperature measurements made using the iButtons to the estimates from the PRISM data described above.

To qualitatively understand how populations’ estimated upper thermal limits compared to source environmental conditions, we also calculated the number of days exceeding 31.6LJ during the adult activity period (*e.g.,* March –July), as this was the estimated upper thermal limit for adult lifespan (the lowest limit for any trait). However, we did not investigate correlations between this environment variable and thermal performance characteristics to minimize multiple testing. The *Ae. sierrensis* dormancy and adult activity windows described above were informed by prior research in this system^39,44^, as well as extensive *Ae. sierrensis* surveillance data available from VectorSurv (https://gateway.vectorsurv.org). Specifically, we examined variation in trapped adult abundance across the year using surveillance data from 2000 – 2020 for the trap closest to each of our collection sites (Supplemental Figure S4).

### Analysis: Fitting thermal response curves

To estimate the thermal limits and performance characteristics of each trait and population, we fit thermal response curves to the experimental data using a Bayesian approach following methods described in detail in Shocket et al. 2020^45^. We first visually inspected the temperature-performance data to determine the most appropriate functional form of the thermal response for each trait. Consistent with prior work, we used quadratic fits truncated to a maximum of 1 for larval and pupal survival, quadratic fits for adult lifespan, and Brière fits for larval and pupal development rate and fitness^45,46^ (Supplemental Table S3).

We fit a first set of Bayesian models for each combination of trait and population across temperatures using uniform priors for the thermal limit parameters bounded by biologically plausible temperature cut-offs as in prior studies^11,45–48^ (*i.e.*, trait performance was set to zero below 0LJ and above 40-45LJ depending on the trait; Supplemental Table S3). For larval and pupal development rate, adult lifespan, and fitness, we modeled the observed data as normally distributed with the mean predicted by the thermal response function at that temperature and the standard deviation, σ, as a gamma distributed parameter, 1/σ^2^, with shape parameter α = 0 and rate parameter b = 1000. For larval and pupal survival probabilities, we modeled the observed data as binomially distributed with the probability and number of trials based on the proportional survival and sample size for that temperature – population combination. We truncated thermal response functions at zero for all traits, as well as at one for survival probability traits. We fit models using Markov Chain Monte Carlo (MCMC) sampling, which uses simulation to approximate the posterior distribution, using the ‘R2jags’ package^49^. For each thermal response, we ran three independent chains with a 5,000-iteration burn-in, and thinned the chains by saving every eighth iteration. This fitting process produced 7,500 values in the posterior distribution for each parameter of the thermal response function (*i.e.*, T_min_, T_max_ and q) and enabled us to calculate additional derived quantities for each trait and population including the maximum trait performance value (P_max_), the temperature at maximum performance (T_opt_), and the temperature range where performance is at least 50% of the maximum (T_breadth_; See Supplemental Figure S5 for theoretical thermal performance curve). We refer to the above fitting process as our ‘low information’ model specification.

To reduce the uncertainty in our parameter estimates, we then fit a second set of models—the main models presented in the text—using informative priors generated using a two-step process. In the first step, we specified low information priors as described above for each population and trait but using only the temperature-performance data from the other nine populations (*i.e.,* a ‘leave-one-out’ approach^45^). We fit a Gamma probability distribution to the posterior distributions of each thermal response parameter using the ‘MASS’ package^50^. We then used these hyperparameters as informative priors in a second round of model fitting. To ensure the hyperparameters did not have an outsized influence on the resulting posterior distributions, we increased the variance of the priors through multiplication by a constant k, set at 0.1 or 0.01, depending on the trait (Supplemental Table S3). The parameter estimates from this ‘informative’ model specification are presented as the main results in the text but did not differ qualitatively from those made through the ‘low information’ model specification presented in the supplement. When investigating variation in thermal performance parameters, we interpreted non-overlapping credible intervals as biologically meaningful and statistically supported differences between populations and/or traits^51–55^. It is worth noting that the leave-one-out informative prior approach biases our thermal performance curve fits to be more similar across populations, making the resulting estimates of differences among populations conservative. On the other hand, this approach has the advantage of realistically constraining uncertainty, for example in cases where a trait was poorly quantified at a given temperature (*i.e.*, few individuals in a given population survived to the relevant life stage).

## Results

### How much does thermal tolerance vary between populations across the species range?

We investigated variation in mosquito thermal performance between 10 populations across the species range. For each population, we characterized the thermal performance of life history traits constituting fitness by fitting thermal response curves (Figure 2, top panel) to our experimental data and estimating the thermal limits and thermal optima (Figure 2, bottom panel).

**Figure 2.**
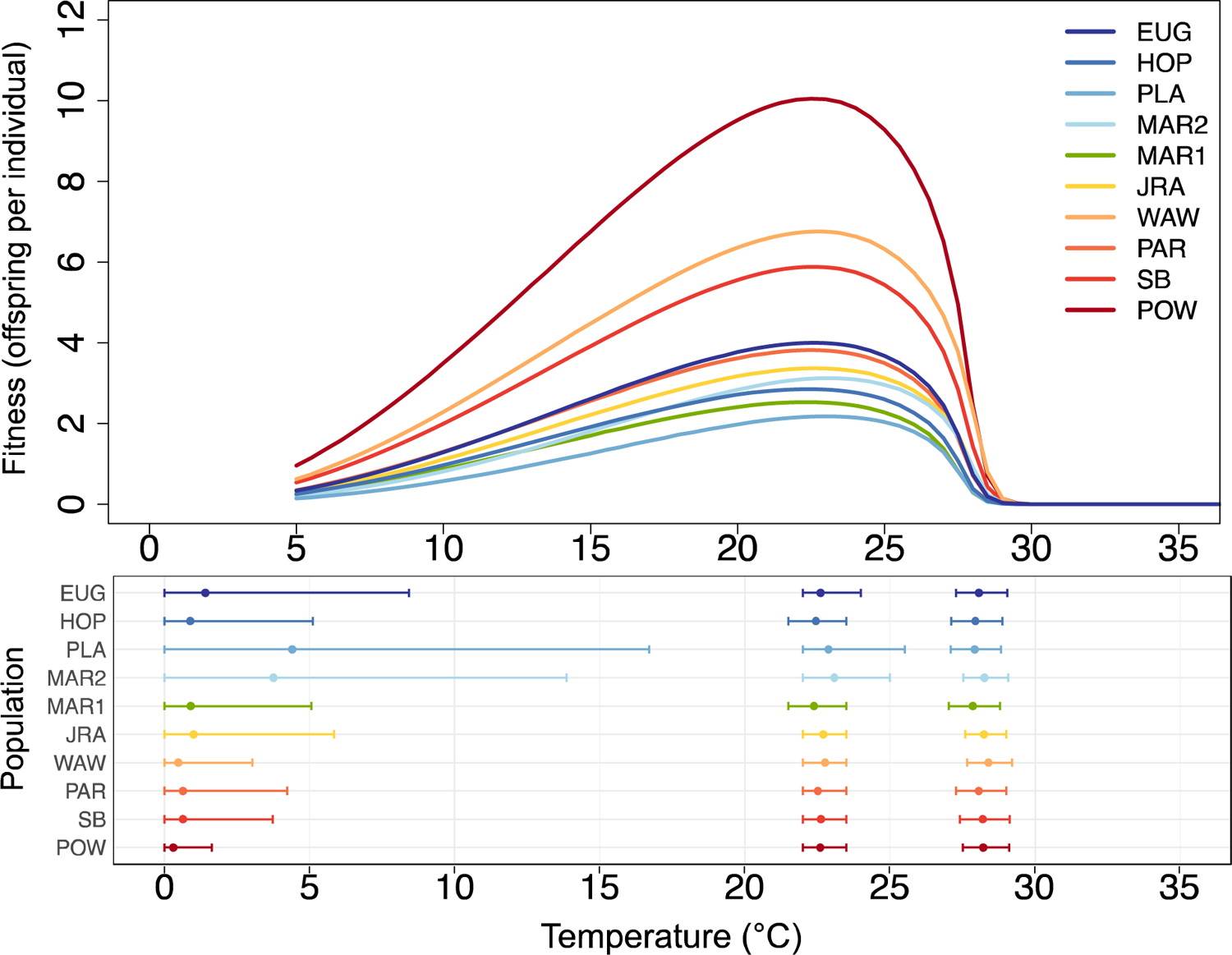
Populations vary minimally in their thermal limits and optima for fitness. In the top panel, each line denotes the mean thermal performance of our fitness proxy for one population. In the bottom panel, points denote estimated thermal performance parameters for our fitness proxy for each population, including lower thermal limit (left), thermal optima (middle), and upper thermal limit (right). Error bars denote the 95% credible intervals for each parameter. In both panels, populations are colored and ordered by their latitude of collection from north (blue) to south (red); this color scheme and ordering is consistent across all figures in the paper.

For our fitness proxy, we found very little variation in thermal tolerance between populations (Figure 2). Specifically, both upper thermal limits and thermal optima varied by <1LJ across all populations, ranging from 27.8 – 28.4LJ and 22.4 – 23.1LJ, respectively. Further, the 95% credible intervals for these parameters overlapped for all populations, indicating non-significant differences between populations. Populations displayed greater, but non-significant, variation in their lower thermal limits for fitness, ranging from 0.3 – 4.6LJ. These results were highly similar when using the low information model specification (Supplemental Figures S8-9). While it was not the focus of this study, we did also find that populations varied in maximum fitness (P_max_)—when averaging across temperature treatments, population’ maximum fitness ranged from an estimated 2.2 – 10.1 offspring per individual (Supplemental Figure S6). We did not detect between-population variation in the thermal breadth of fitness (Supplemental Figure S6), nor any consistent correlations between fitness thermal performance characteristics (*i.e.*, between P_max_ and T_breadth_ or between P_max_ and T_opt_) among populations. These analyses and results are discussed further in the Supplemental Results.

As with fitness, we found minimal variation in thermal tolerance between populations for most individual life history traits (Figures 3-4). In particular, for all life history traits, both upper and lower thermal limits varied by <3°C across populations (Figure 4, Supplemental Figure S17). Similarly, thermal optima varied by <1.5°C for all traits except larval and pupal survival, for which our estimates had the greatest uncertainty (partly due to high juvenile survivorship across the intermediate temperature treatments). Variation between populations was non-significant (*i.e.,* 95% credible intervals overlapped for all populations) for nearly all life history traits and thermal performance parameters, with three exceptions: the upper thermal limits (T_max_) of larval and pupal development rates, and the thermal optima (T_opt_) of pupal development rates. Upper thermal limits for larval and pupal development rates each varied by 1.6°C across populations (33.3 – 34.9°C and 32.1 – 33.7°C, respectively), while the thermal optima of pupal development rate varied by 1.4°C (26.3 – 27.7°C).

**Figure 3.**
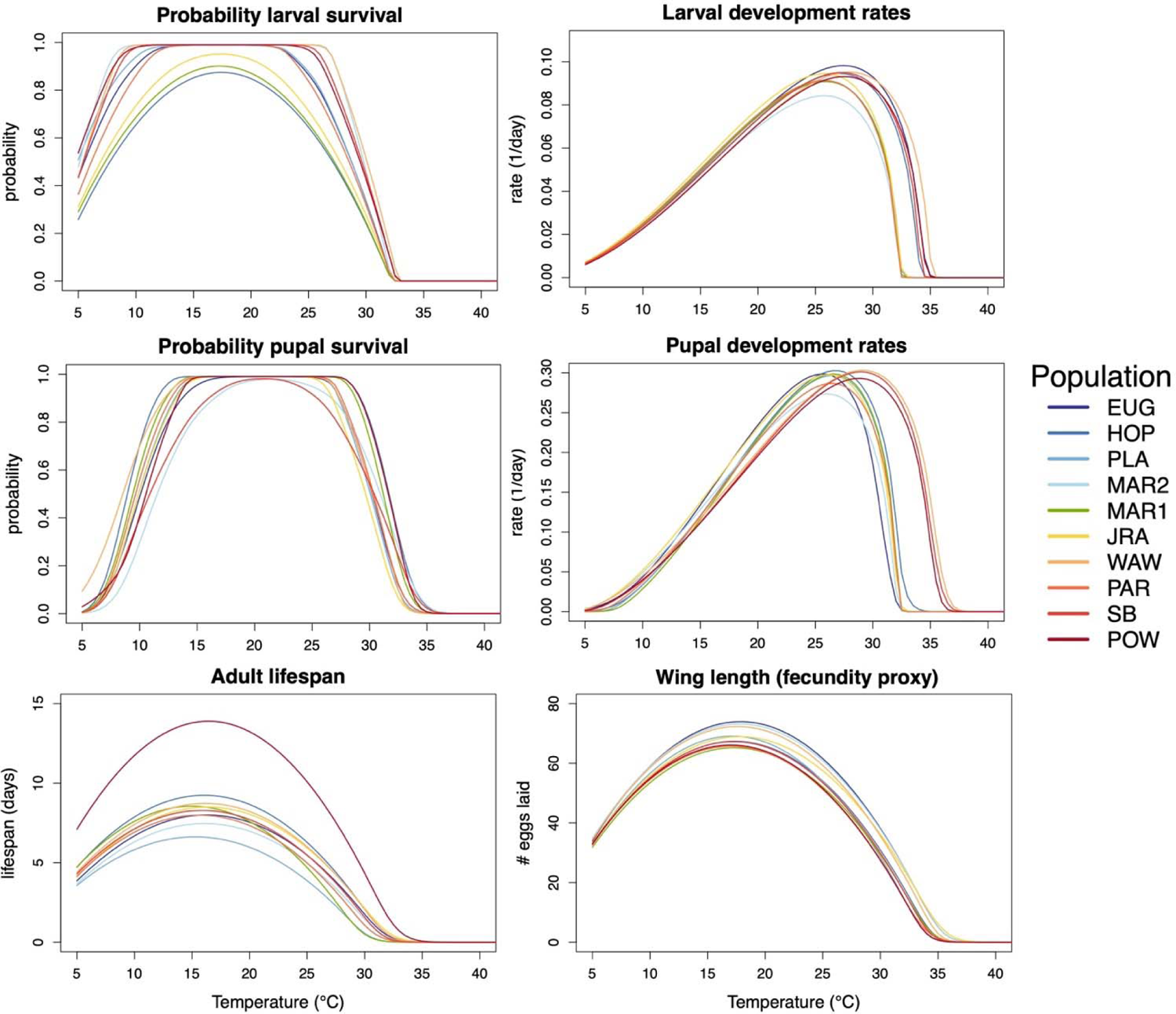
For most life history traits, thermal performance varies minimally between populations. Each curve denotes the average thermal performance for one population for a given trait. Populations are colored and ordered in the legend by their latitude of collection.

**Figure 4.**
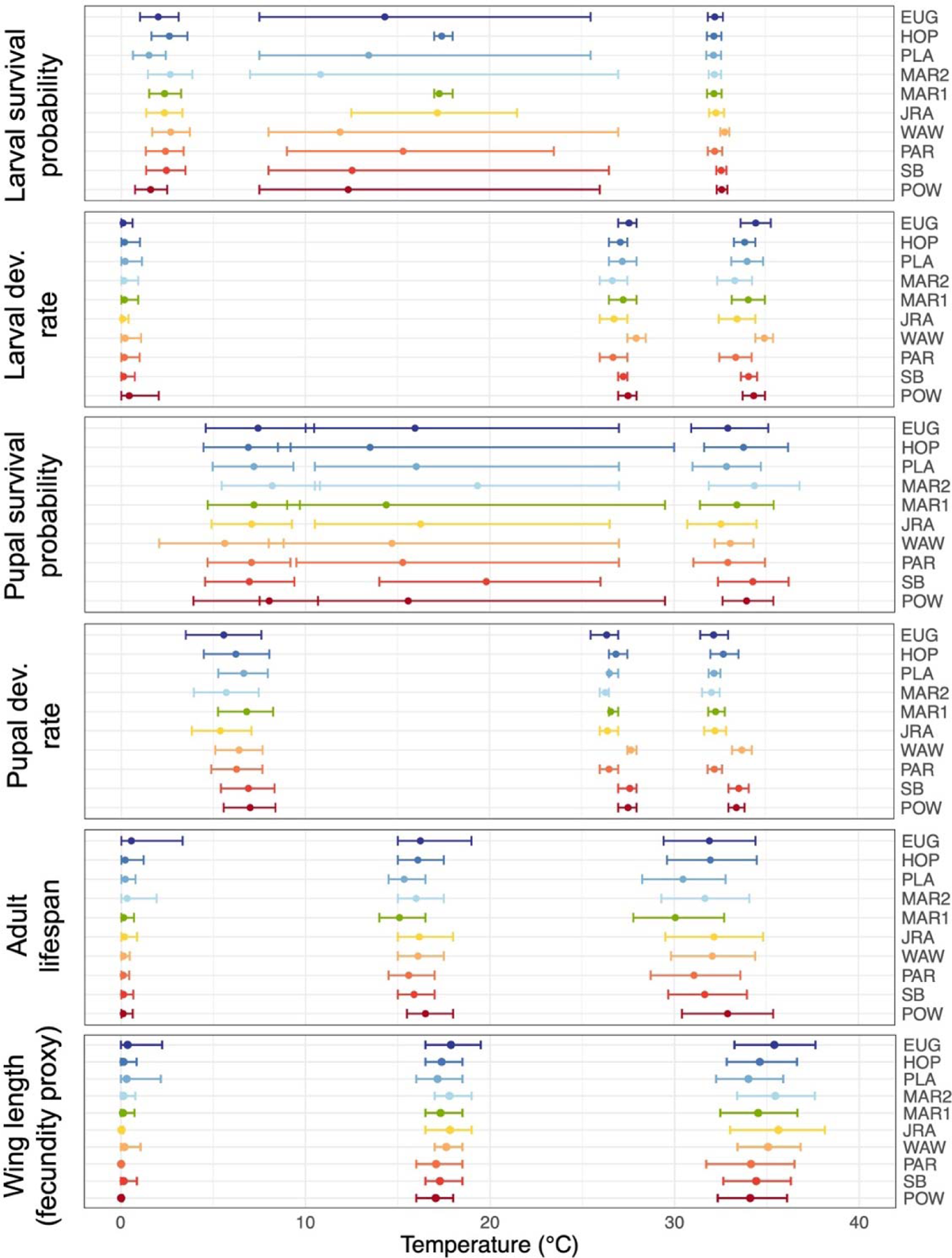
Thermal limits and optima vary between life history traits, but minimally between populations. Lower thermal limits, thermal optima, and upper thermal limits for each life history trait and population (left, middle, and right points and error bars in each panel, respectively). Thermal performance parameter estimates are derived from the thermal performance curves for traits for which the means are depicted in Figure 3. Points and error bars denote the mean and 95% credible intervals for each parameter, respectively. Populations (listed on the right) are colored and ordered by latitude of collection. Units of development rates and lifespan are 1/days and days, respectively. Note that survival probability curves that are truncated at one have very uncertain optimal temperatures because a wide range of temperatures have similarly high survival probability.

### Is variation in thermal performance correlated with the source thermal environment?

To assess evidence of local thermal adaptation, we investigated the relationship between the source thermal environment (Table 1) and experimentally measured thermal performance parameters, using only the parameters with biologically significant between-population variation (*i.e.*, those where populations had non-overlapping 95% credible intervals). This included the upper thermal limits (T_max_) of larval and pupal development rates, and the thermal optima (T_opt_) of pupal development rates.

**Table 1.** Thermal characteristics of the source environment for each population listed in order of decreasing latitude (*i.e.*, north to south). Values represent averages from 2000 – 2020, calculated from PRISM climate data at a 1 km buffer around the sampled tree hole. Seasonal temperature variation is defined as the difference between the mean warmest month temperature and the mean coolest month temperature. Warm-season maximum is defined as the mean daily maxima in the Spring and Summer. The # days > 35 or 31.6°C refer to the average number of days where the maximum temperatures exceeded the stated threshold, either across the year, or when considering only non-dormant periods (January – July) or adult activity periods (March – July). See Supplemental Figure S18 for correlations between temperatures variables and Supplemental Figure S19 for comparisons between the PRISM and iButton temperature estimates for the ‘SB’ and ‘POW’ populations.

| Pop. | Annual mean temp. (°C) | Jan – March mean temp. (°C) | Seasonal temp. variation (°C) | Warm-season maximum (°C) | # days > 35°C |  | # days > 31.6°C |  |
| --- | --- | --- | --- | --- | --- | --- | --- | --- |
|  |  |  |  |  | Across year | Jan. - July | Across year | March - July |
| EUG | 11.48 | 6.33 | 16.01 | 22.69 | 4.29 | 2.23 | 18.33 | 9.45 |
| HOP | 14.57 | 9.41 | 15.38 | 26.77 | 24.90 | 11.95 | 57.67 | 28.863 |
| PLA | 16.08 | 9.91 | 18.08 | 28.04 | 29.95 | 17.00 | 76.24 | 40.27 |
| MAR2 | 14.24 | 10.31 | 11.62 | 24.77 | 8.71 | 4.18 | 25.95 | 12.81 |
| MAR1 | 14.04 | 10.18 | 11.29 | 24.41 | 6.33 | 3.05 | 20.86 | 10.36 |
| JRA | 15.45 | 11.49 | 11.61 | 25.29 | 10.43 | 4.77 | 27.14 | 13.54 |
| WAW | 15.83 | 8.73 | 20.51 | 28.19 | 41.57 | 20.95 | 87.76 | 44.18 |
| PAR | 14.40 | 9.88 | 13.26 | 26.52 | 14.71 | 5.77 | 46.67 | 22.73 |
| SB | 16.44 | 11.82 | 13.51 | 27.86 | 26.81 | 11.77 | 59.95 | 29.72 |
| POW | 18.75 | 15.08 | 11.65 | 27.77 | 18.14 | 5.09 | 50.62 | 21.09 |

We found several correlations that reflected patterns of local thermal adaptation. In particular, we found that T_max_ and T_opt_ of pupal development were positively correlated with annual mean temperature, maximum daily temperatures in the Spring and Summer, and the number of days exceeding 35°C (r: 0.64 – 0.71; Figure 5). Together, this is consistent with local thermal adaptation of pupal development rate to high temperatures. By contrast, T_max_ of larval development rate was not strongly correlated with any source temperature variable. We note that these reported correlations are only statistically significant (p < 0.05) prior to adjustment for multiple comparisons, the necessity of which is debated when making only specific, biologically meaningful comparisons (as we have done here) rather than all possible comparisons^58,59^. The majority of the above correlations remained significant after removing ‘POW’ (Supplemental Table S6), the lowest latitude population, indicating that our findings of thermal adaptation are not solely driven by this population.

**Figure 5.**
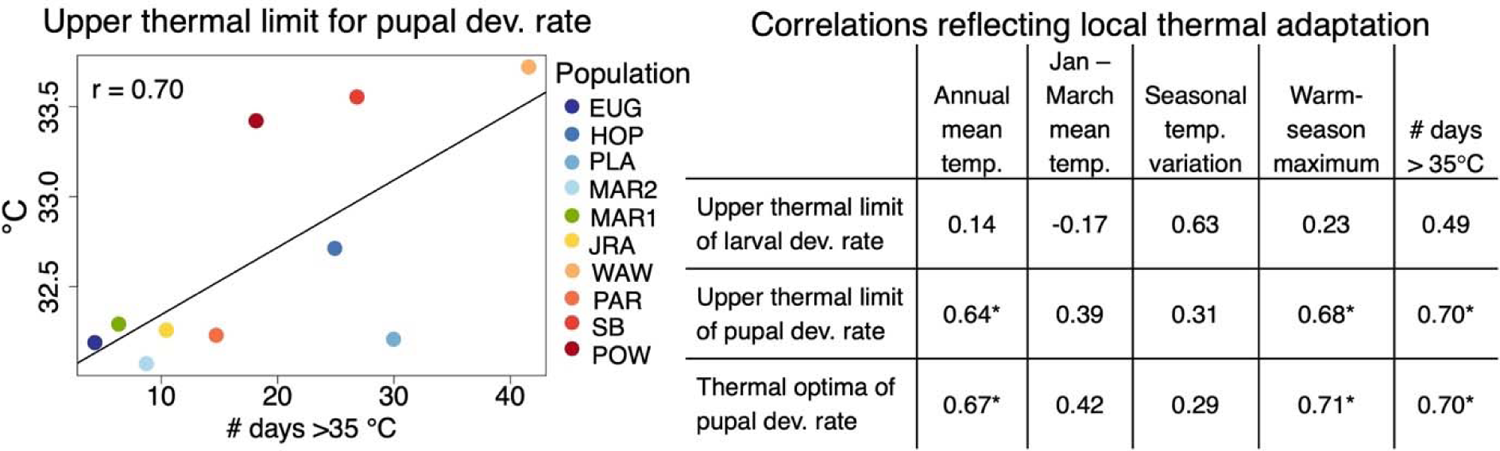
Evidence of local thermal adaptation. Correlations between the source thermal environment and population thermal performance provide evidence of local thermal adaptation (right). Statistically significant Pearson’s correlations (r; p < 0.05) are denoted with (*). Note that correlations were only examined for traits with significant-between population variation. The relationship between upper thermal limits for pupal development rate and the number of days with temperatures exceeding 35°C (one of the significant correlations noted in the table) is visualized in the plot on the left.

We also found that maximum fitness (P_max_), which varied significantly between populations, was positively correlated with annual mean temperature (r = 0.66; no correlations with other temperature variables were statistically significant). This result that populations from warmer climates generally have higher maximum fitness has frequently been found in other ectotherms^60–62^, but does not necessarily reflect local thermal adaptation, in which peak fitness is expected to occur at conditions most similar to the source environment.

### At present, how often do environmental temperatures exceed mosquito populations’ upper thermal limits?

We found that for all populations, temperatures in the surrounding environment already exceed our estimated upper thermal limits. In particular, the number of days per year with temperatures exceeding 35°C—above the highest upper thermal limit we estimated for any life history trait— ranged from 2 to 20 days (Table 1). This metric specifically excluded times of the year when *Ae. sierrensis* populations are likely in dormancy (*e.g.,* August – October) – if all months were included, there were an average of 4 to 42 days exceeding this threshold. Similarly, the number of days exceeding 31.6°C—the lowest estimated upper thermal limit (adult lifespan)—ranged from 9 to 40 days during adult activity season (*e.g.,* March – July) or 18 to 88 days across the entire year.

The above estimates are based on PRISM climate data, which captures air temperature in the broader surrounding environment, but not necessarily the precise temperature experienced in a given tree hole. For two populations, we were able to record temperatures within the tree hole for approximately one year following larval collection. We found that these direct measurements were strongly correlated with temperature estimates from the PRISM climate data (r = 0.91, 0.87 for daily temperature estimates for the SB and POW populations, respectively; Supplemental Figure S19). For these populations, the iButton recorded daily temperatures that were, on average, 0.70°C higher (SB) or 3.0°C lower (POW) than the PRISM estimates. In both locations, tree hole temperatures exceeded 31.6°C on several days (Supplemental Figure S19), indicating that populations are exposed to temperatures above their estimated upper thermal limits for adult lifespan even within this microhabitat.

## Discussion

In one of the largest-ranging studies of standing variation in mosquito thermal tolerance to date, we found limited evidence of variation between populations in the thermal responses of fitness and life history traits. Specifically, in our common garden experiment using ten *Aedes sierrensis* populations spanning over 1,200 km, we found the upper thermal limits and thermal optima for fitness each varied by <1LJ across all populations (27.8 – 28.4LJ and 22.4 – 23.1LJ, respectively; Figure 2). This level of variation in upper thermal limits across latitude (*i.e.*, 0.6°C across populations spanning 10° of latitude) is large relative to previous studies in terrestrial ectotherms (0.3°C per 10° latitude^57^); however, it is considerably less than the level of variation in environmental temperature across this range, and likely less than the extent of warming expected in this region in coming decades^63^.

Our finding of minimal variation in mosquito thermal tolerance across the species range is consistent with prior findings in a broad range of ectotherm species^64,65^. For taxa including insects, arachnids, reptiles, and amphibians, upper thermal limits typically vary little across wide climatic and latitudinal gradients^57,65–67^, a pattern that has been suggested to reflect hard evolutionary constraints on heat tolerance^68,69^. Although the underlying mechanism remains unclear, the evolution of heat tolerance may be limited by genetic constraints (*e.g.,* low heritability) and/or biochemical constraints (*e.g*., limits on enzyme stability at high temperatures)^64,70,71^. Alternatively, this pattern could be driven by behavioral strategies enabling populations to experience and adapt to similar thermal regimes across their range^72^, and/or trade-offs in adapting to temperature versus other abiotic or biotic selection pressures^73^.

Despite generally limited variation in thermal tolerance between populations, we did observe meaningful variation in the thermal responses of larval and pupal development rates (Figure 3). For both traits, upper thermal limits varied significantly, and by approximately 1.6°C across populations—over twice as large as the variation estimated in fitness upper thermal limits in our study and five times the average across terrestrial ectotherms spanning a similar latitudinal extent^57^ (Supplemental Table S4). Further, for pupal development rate, we found that variation in populations’ thermal optima and upper thermal limits was strongly correlated with variation in the source thermal environment. Specifically, populations from environments with higher mean and extreme temperatures had higher thermal optima and limits for pupal development rate than those from cooler source environments, providing clear evidence for local thermal adaptation in this trait (Figure 5).

That thermal adaptation was observed specifically in pupal development rate may be due to the seasonal ecology of *Ae. sierrensis* making the pupal life stage the most vulnerable to high temperatures. In particular, *Ae. sierrensis* eggs and larvae undergo a period of dormancy and are primarily active earlier in the season, which may buffer these life stages from high temperature extremes, while adults may avoid high temperatures through movement to cooler microhabitats^24,39^. Conversely, pupae have limited capacity for movement, no period of dormancy, and typically begin development in the spring, which can have highly variable thermal conditions across years and include high temperature extremes. This life history trait may thus experience the strongest thermal selection pressure given the exposure to thermal stress and a lack of other coping strategies. By measuring the thermal performance of traits across the species life cycle, and using many populations from across a wide thermal gradient, we were able to detect this specific evidence of thermal adaptation, which has not been clearly identified in prior investigations of thermal adaptation in other mosquito species^17,18,20^.

Despite this evidence of local thermal adaptation, the potential for further evolutionary adaptation to warming could be limited. In addition to the minimal variation observed in upper thermal limits for most traits, we found that temperatures at all source environments already exceed our estimated upper thermal limits (Table 1). In particular, environmental temperatures at each of our collection sites were at or above 35°C—exceeding the highest upper thermal limit we estimated for any trait—for an average of 2 to 20 days out of the potential *Ae. sierrensis* activity season (January - July). Similarly, environmental temperatures exceeded 31.6°C—the lowest upper thermal limit across measured life history traits (adult lifespan, Figure 4)—for 9 to 40 days during this period. Thus, populations may already be exposed to temperatures beyond their estimated upper thermal limits; however, the extent to which this indicates climate vulnerability depends on the time scales over which these high temperatures occur. In particular, short-term thermal extremes (*e.g.,* one to several hours) that are followed by cooler temperatures could be tolerated through heat stress repair, as has been found to occur during night-time in other ectotherm species^74^. As our experiments involved constant-temperature exposure, we were unable to test whether such repair mechanisms could enable higher thermal tolerance – incorporating diurnal temperature variation is an important next step for future experiments. In addition to short-term heat repair, other strategies besides evolutionary adaptation, such as seasonal life cycles and microhabitat selection may be important for sustaining *Ae. sierrensis* under rapid climate warming. Accordingly, the majority of days exceeding the 35°C and 31.6°C thresholds at our collection sites occurred after July, when most individuals in the population are likely in the dormant egg stage (Supplementary Figure S3). Further, the tree hole microhabitat in which *Ae. sierrensis* completes most of its life cycle may be cooler than the surrounding environment, further buffering individuals from thermal extremes (although we found this was not consistently the case; Supplementary Figure S18).

Phenological and behavioral strategies for mitigating thermal danger may be similarly important for other mosquito and ectotherm species to persist under ongoing climate warming^72,75,76^. For example, *An. gambiae* in the Sahel have been shown to persist during the arid summers by entering a prolonged period of dormancy^77^, and winter dormancy responses are widespread among mosquito species, likely facilitating their geographic expansion^78,79^.

Similarly, biting activity in several mosquito species has been found to shift during warmer months from dusk to late at night, although this was not conclusively linked to temperature. Behavioral avoidance of high temperatures (typically >30°C) has been documented in adult *Aedes, Anopheles*, and *Culex* spp. under lab conditions^82–84^, and some evidence of preference for cooler, shaded oviposition sites in warm climates has been found in field settings^85,86^. These types of strategies can have a large impact on buffering individuals from thermal stress^72,75,87^, but may dampen selection for greater thermal tolerance, further decreasing the likelihood of evolutionary adaptation (termed the ‘Bogert effect’^88^). Identifying the extent of behavioral thermoregulation and temperature-driven changes in phenology in natural settings, and their potential to enable mosquito persistence under climate warming are important directions for future research.

Our experiment focused on the impacts of constant temperatures on mosquito trait performance—an important first step in characterizing thermal tolerance for a given species. However, changes in temperature fluctuations and short-term thermal extremes are key components of climate warming projections and can have a large impact on mosquito life histories^89–94^. In particular, coping with large fluctuations in temperature and/or acute thermal extremes may require a different set of physiological or behavioral strategies than coping with constant warm temperatures^95–97^. Thus, patterns of mosquito thermal adaptation to these aspects of temperature could differ from those estimated here. However, to our knowledge, no studies have yet measured variation in mosquito responses to mean, fluctuating, and extreme temperatures between populations. Prior studies in other ectotherm species have tested whether thermal performance under fluctuating temperatures can be predicted qualitatively from thermal performance curves estimated at constant temperatures, finding mixed results^98,99^. Experimentally testing this approach in mosquitoes and estimating mosquito performance under thermal regimes that reflect natural conditions using populations from across the species range are important future directions.

## Authors’ Contributions

L.I.C. and E.A.M. conceived of and designed the project. L.I.C., J.E.F, and K.P.L. performed the field collection and laboratory rearing. L.I.C., J.E.F., K.P.L., and A.S.L. conducted the experiment. L.I.C. conducted the analyses and drafted the manuscript. All authors revised the manuscript and read and approved the final version.

## Acknowledgements

We received tremendous support with field collection and rearing from many vector control officials including Bret Barner, Peter Bonkrude, Joel Buettner, Angela Caranci, Kelly Liebman, Nathan McConnell, Angie Nakano, Andrew Rivera, Karen Schultz, Mary Sorenson, and Greg Williams, and Jasper Ridge staff scientist Nona Chiariello. We acknowledge Jasper Ridge Biological Preserve and Hopland Research and Extension Center as valuable field collection sites. We are grateful to Isabel Delwel, Dylan Loth, Desire Uwera Nalukwago, and Mallory Harris for help setting up the experiment, and the Mordecai lab members broadly for feedback on experimental design and the manuscript. We thank Marta Shocket for sharing code that helped with the Bayesian modeling approach. This work was supported by the Philippe Cohen Graduate Fellowship, the Lewis and Clark Field scholarship, the Pacific Southwest Center of Excellence in Vector-Borne Diseases Training Grant, the Stanford Center for Computational, Evolutionary and Human Genomics, the Stanford Center for Innovation in Global Health, Stanford’s Woods Institute for the Environment, the Bing-Mooney Fellowship, the National Science Foundation Postdoctoral Research Fellowships in Biology Program (2208947), the National Science Foundation (DEB-2011147, with Fogarty International Center), and the National Institutes of Health (R35GM133439, R01AI168097, and R01AI102918).

## Conflict of Interest

The authors declare that they have no conflicts of interest.

## Data accessibility

All data and code used in this project are publicly available on GitHub: https://github.com/lcouper/MosquitoThermalAdaptation

## Notes

https://github.com/lcouper/MosquitoThermalAdaptation

### Competing Interest Statement

The authors have declared no competing interest.

### Summary of Updates

– clarification around the wing length measurement as a proxy for fecundity – additional details have been added regarding methodological choices (e.g., bounding of uniform priors, correlational analyses, interpretation of non-overlapping credible intervals) – clarification that we did not eliminate cross-generational effects – updates to figures to display temperature variation across the collection sites – reduced emphasis on climate vulnerability and increased discussion about the time scales of thermal extremes

