## Supplemental Information for "Mosquito thermal tolerance is remarkably constrained across a large climatic range"

**This file includes:**

Supplemental Methods

Supplemental Results

Supplemental Tables S1 – S6

Supplemental Figures S1 - S19

References

**Supplemental Methods**

*Validation experiment: Assessing wing length as a proxy for fecundity*

To validate the Washburn et al. (1989)^1^ relationship between wing length and fecundity, we reared a separate, smaller number of individuals from each of the 10 focal populations (excluding MAR2 due to insufficient remaining eggs). In this validation experiment, all individuals were reared at 17℃ until adulthood. Exactly one week after adult emergence, individual adult females were offered a 30-minute blood meal consisting of defibrinated sheep’s blood supplied through a Hemotek (Blackburn, UK) membrane feeder. Any blood-fed individual was moved to an individual 12 oz plastic container with an oviposition cup and one 10% sugar-soaked cotton ball. Individuals from each population were then assigned to remain at either 13℃ (n = 24 total) or 24℃ (n = 25 total) for the remainder of the experiment, to assess if the estimated wing length-fecundity relationship held across temperatures.

Each day of the experiment, we checked the containers for the presence of eggs, resupplying water and/or sugar as needed, and noting any adult deaths. On the first day that we detected eggs for an individual, we removed the oviposition cup, and flash froze the adult by placement at -80℃. We then counted the number of eggs laid via microscopy and measured the left wing of each mosquito as before. At both 13℃ and 24℃, we found a highly similar relationship between wing length and fecundity as Washburn et al. 1989 (Supplemental Figure S2). Thus, we proceeded to use the Washburn et al. 1989 relationship to estimate fecundity from the adults in our experiment as it had a larger sample size.

*Determining age at reproductive maturity*

To determine the age of reproductive maturity, as used in our approximation of individual-level fitness, we first measured the minimum number of days from adult eclosion to egg-laying at a given temperature, as observed in the validation experiment described above. That is, we offered a blood meal one week after adult emergence, then held individuals at different constant temperatures and recorded the minimum number of days until egg-laying. In addition to the 13 and 24℃ temperature treatments used to validate the wing length-fecundity relationship, for this measurement, we also held individuals at 17 (n= 27) and 28℃ (n =28), to capture the range of temperature treatments used in the main experiment. In our pilot experiments, no individuals survived long enough to take a blood meal at 32℃ and no individuals laid eggs at 5℃, therefore we did not include these temperature treatments here. We found the minimum number of days from blood-feeding to egg-laying were: 13 days at 13℃, 7 days at 17℃, and 6 days at 24 and 28℃. We then determined the age at reproductive maturity as this time plus four days – the minimum time from eclosion to blood-feeding, which we have observed consistently across the 13 - 28℃ temperature treatments used here.

**Supplemental Results**

*Variation in thermal performance between traits*

Thermal limits and optima displayed much greater variation between traits than between populations (Figure 3-4, Supplemental Figures S11-17; Supplemental Tables S4-5). Averaging across populations, the highest upper thermal limits occurred for estimated fecundity (34.7°C) and larval development rate (34.0°C) and the lowest upper thermal limits occurred for adult lifespan (31.6°C). Larval development rate also had the highest thermal optimum (27.2°C), while larval survival had the lowest (14.3°C). When averaging across all populations, we found that thermal optima and lower thermal limits varied more between traits than did upper thermal limits (Supplemental Tables S4-5)—a finding that is consistent with patterns of thermotolerance in ectotherms^2,3^ and may reflect the presence of shared physiological constraints on upper heat tolerance across traits.

*Additional thermal performance characteristics: maximum fitness and* *thermal breadth*

In addition to estimating thermal limits and optima, our Bayesian approach for fitting thermal performance curves enabled us to estimate maximum trait performance ($P_{\max}$), and thermal breadth of performance $(T_{\mathrm{breadth}}$) for each population. While it was not the focus of this study, we estimated these additional thermal performance characteristics for fitness for each population.

We found that populations varied substantially in maximum fitness performance (Figure 2; Supplemental Figures S8-9) – averaging across temperature treatments, populations’ maximum fitness ranged from an estimated 2.2 – 10.1 offspring per individual. Populations with higher maximum fitness were typically those from lower latitudes, with the population from the lowest latitude (‘POW’) achieving the highest maximum fitness across all temperatures (Figure 2, Supplemental Figures S7-8). In particular, this population had the highest proportion of individuals reaching adulthood and surviving to the date of potential oviposition (*i.e.*, achieving non-zero fitness, Supplemental Figure S10), which was the major bottleneck in this experiment: only 141 out of the 1,736 total individuals (8%) used in the experiment achieved non-zero fitness. This low-latitude population also had the highest estimated fecundity of the surviving adults at nearly all temperatures (Supplemental Figure S10), suggesting that it did not face trade-offs between survival and reproduction in this experimental setting. Unlike maximum fitness performance, we found populations did not vary significantly in their thermal breadth (but ranged from 12.9 – 14.8℃; Supplementary Figure S6).

*Correlations in thermal performance characteristics*

While it was not the focus of this study, by estimating multiple parameters of thermal performance (*i.e*., $T_{\max}$, $T_{\min}$, $T_{\mathrm{opt}}$, $P_{\max}$, and $T_{\mathrm{breadth}}$) for each population, we could investigate relationships between these parameters and evaluate evidence for different hypotheses of thermal performance curve evolution. The ‘specialist-generalist hypothesis’ posits that high performance at one temperature comes at the cost of reduced performance breadth due to trade-offs in enzyme stability and flexibility^4^. From our experimental data, we found that higher maximum fitness was not associated with reduced thermal breadth. In particular, the population with the highest maximum performance (‘POW’) also had the widest thermal breadth, while the population with the lowest maximum fitness (‘PLA’) had the narrowest thermal breadth (main text Figures 2-3). Maximum performance and thermal breadth were weakly positively correlated across populations, but the strength varied when using informative versus low information priors (r = 0.16, 0.04, respectively), suggesting that the relationship was influenced by the precise shape of the fitted thermal response curve. Next, we investigated support for the ‘hotter-is-better’ hypothesis, which posits that genotypes with higher thermal optima have higher trait performance than those with cooler thermal optima because warmer temperatures accelerate enzymatic rates and metabolism^5^. We did not detect a consistent association between maximum fitness performance and thermal optima in our populations—the correlation was weakly positive when using informative priors (r = 0.17), and negative when using low information priors (r = -0.75).

*Source thermal environments*

Field-collected mosquito populations originated from a wide range of thermal environments, with average annual mean temperatures ranging from 11.5 – 18.7 °C, seasonal temperature variation ranging from 11.3 – 20.5°C, and daily maxima in the Spring and Summer ranging from 22.7 – 28.2°C (Table 1). In general, source environment mean temperatures were positively correlated with high temperature extremes and negatively correlated with seasonal temperature variation, but the strength of the correlations varied between 0.06 – 0.87 (Supplemental Figure S18), indicating that no single temperature variable holistically characterizes the source thermal environment.

**Supplemental Table S1.** Population collection metadata including population abbreviation, site name, collection date, latitude, longitude, elevation, and tree species.

| **Pop.** | **Site name** | **Collection date** | **Latitude** | **Longitude** | **Elevation (m)** | **Tree species** |
| --- | --- | --- | --- | --- | --- | --- |
| EUG | Eugene | 4/14/22 | 44.023204 | -123.098694 | 178 | Unknown oak |
| HOP | Hopland reserve | 10/31/21 | 39.0099819 | -123.0722175 | 449 | Deciduous oak |
| PLA | Placer county | 3/29/22 | 38.9039747 | -121.0543602 | 330 | Unknown oak |
| MAR2 | Olompali | 11/1/21 | 38.1525295 | -122.5708248 | 21 | Live oak |
| MAR1 | Rush creek | 11/1/21 | 38.1304913 | -122.543991 | 30 | Live oak |
| JRA | Jasper Ridge | 11/11/21 | 37.40805 | -122.23703 | 145 | Live oak |
| WAW | Oakhurst | 4/3/22 | 37.391291 | -119.633103 | 1023 | Unknown oak |
| PAR | Paso robles | 2/22/22 | 35.455543 | -120.688386 | 308 | Unknown oak |
| SB | Paradise campground | 2/17/22 | 34.54276 | -119.81219 | 280 | Coast live oak |
| POW | Powder canyon | 1/26/22 | 33.9638888 | -117.9243188 | 229 | Coast live oak |

**Supplemental Table S2.** Number of *Ae. sierrensis* individuals randomly assigned to each temperature treatment. Note that for all populations, more individuals were assigned to the 24, 28, and 32°C treatments as higher juvenile mortality was expected at these temperatures based on pilot experiments.

|  | **Temperature treatment** | | | | | |
| --- | --- | --- | --- | --- | --- | --- |
| **Population** | **5°C** | **13°C** | **17°C** | **24°C** | **28°C** | **32°C** |
| EUG | 11 | 10 | 10 | 19 | 31 | 40 |
| HOP | 20 | 15 | 20 | 36 | 40 | 56 |
| PLA | 21 | 17 | 20 | 35 | 43 | 50 |
| MAR2 | 11 | 8 | 9 | 29 | 25 | 50 |
| MAR1 | 21 | 18 | 19 | 37 | 36 | 50 |
| JRA | 21 | 16 | 19 | 34 | 39 | 50 |
| WAW | 20 | 19 | 19 | 37 | 38 | 52 |
| PAR | 20 | 19 | 19 | 36 | 35 | 49 |
| SB | 21 | 17 | 18 | 36 | 37 | 71 |
| POW | 21 | 19 | 19 | 38 | 39 | 51 |

**Supplemental Table S3.** Bayesian model specification for each life history trait. U refers to the uniform distribution. (k) refers to the weighting applied to the informative priors. For the trait functional form, we fit either quadratic ($-q\left( T- T_{o} \right)\left( T- T_{m} \right))$ or Brière or ($\mathrm{qT}\left( T-T_{o} \right)\sqrt{\left( T_{m}-T \right)}$) functions. Under additional trait details, we indicate the populations and traits for which a ‘0’ was manually encoded at 32℃. This was done if the trait could not be empirically measured for that population because no individuals survived to that stage (*e.g*., if all EUG larvae died at 32℃, a ‘0’ would be added for larval development rate at 32℃).

| **Trait (units)** | **Functional form** | **Likelihood distribution** | **Low information prior distribution** | **(k)** | **Additional trait details** |
| --- | --- | --- | --- | --- | --- |
| Larval survival (0/1) | Quadratic (truncated at 1) | Binomial | $T_{0}$ ~ U (0, 24)  $T_{m}$ ~ U (25, 40)  $q$ ~ U (0, 1) | 0.1 | We coded individuals as ‘1’ for the 5℃ treatment if they were still alive on the end date of the experiment (9/28/22) |
| Larval development rate (1/days) | Brière | Normal | $T_{0}$ ~ U (0, 20)  $T_{m}$ ~ U (28, 35)  $q$ ~ U (0, 1) | 0.1 | We manually added a single ‘0’ at 32℃ for populations with no larvae surviving this treatment (MAR1, MAR2, EUG, PLA, JRA, PAR) |
| Pupal survival (0/1) | Quadratic (truncated at 1) | Binomial | $T_{0}$ ~ U (0, 24)  $T_{m}$ ~ U (25, 42)  $q$ ~ U (0, 1) | 0.1 | We manually added a single ‘0’ at 32℃ for populations with no larvae surviving this treatment (MAR1, MAR2, EUG, PLA, JRA, PAR) |
| Pupal development rate (1/days) | Brière | Normal | $T_{0}$ ~ U (0, 20)  $T_{m}$ ~ U (28, 39)  $q$ ~ U (0, 1) | 0.01 | We manually added a single ‘0’ at 32℃ for populations with no pupae surviving this treatment (HOP, MAR1, MAR2, EUG, PLA, JRA, PAR) |
| Adult lifespan (days) | Quadratic | Normal | $T_{0}$ ~ U (0, 24)  $T_{m}$ ~ U (25, 45)  $q$ ~ U (0, 1) | 0.1 | We manually added a single ‘0’ at 32℃ for populations with no adults eclosing at this treatment (HOP, MAR1, MAR2, EUG, PLA, JRA, PAR) |
| Fecundity (estimated # of eggs) | Brière | Normal | $T_{0}$ ~ U (0, 24)  $T_{m}$ ~ U (25, 45)  $q$ ~ U (0, 1) | 0.1 | We manually added a single ‘0’ at 32℃ for populations with no adults eclosing at this treatment (HOP, MAR1, MAR2, EUG, PLA, JRA, PAR) |
| Fitness | Brière | Normal | $T_{0}$ ~ U (0, 20)  $T_{m}$ ~ U (28, 40)  $q$ ~ U (0, 1) | 0.01 |  |

**Supplemental Table S4.** Variation in fitted upper ($CT_{\max}$) and lower ($\mathrm{CT}_{\min}$) thermal limits and thermal optima ($T_{\mathrm{opt}}$) between life history traits. Values denote the mean, and minimum and maximum (in brackets) across all populations for a given trait and parameter.

|  | $\mathbf{CT}_{\mathbf{min}}$ **(°C)** | $\mathbf{T}_{\mathbf{opt}}$ **(°C)** | $\mathbf{C}\mathbf{T}_{\mathbf{max}}$ **(°C)** |
| --- | --- | --- | --- |
| Larval survival | 2.26 [1.50 – 2.69] | 14.25 [10.83 – 17.41] | 32.37 [32.18 – 32.79] |
| Larval development rate | 0.18 [0.07 – 0.43] | 27.22 [26.68 – 27.98] | 34.00 [33.34 – 34.94] |
| Pupal survival | 7.16 [5.61 – 8.18] | 16.07 [13.50 – 19.91] | 33.40 [32.54 – 34.35] |
| Pupal development rate | 6.29 [5.38 – 7.00] | 26.84 [26.31 – 27.69] | 32.67 [32.07 – 33.72] |
| Adult lifespan | 0.20 [0.10 – 0.54] | 15.90 [15.09 – 16.50] | 31.60 [30.06 – 32.90] |
| Fecundity | 0.15 [0.01 – 0.36] | 17.44 [17.05 – 17.88] | 34.73 [34.00 – 35.62] |
| Fitness | 1.54 [0.28 – 4.56] | 22.66 [22.36 – 23.11] | 28.10 [27.81 – 28.39] |

**Supplemental Table S5.** Variation in fitted thermal performance parameters estimated using low information priors. Values denote the mean, and minimum and maximum (in brackets) across all populations for a given trait and parameter. See Supplemental Table S4 for corresponding parameters fit with informative priors.

|  | $\mathbf{CT}_{\mathbf{min}}$ **(°C)** | $\mathbf{T}_{\mathbf{opt}}$ **(°C)** | $\mathbf{C}\mathbf{T}_{\mathbf{max}}$ **(°C)** |
| --- | --- | --- | --- |
| Larval survival | 1.67 [0.40 – 5.04] | 11.44 [5.38 – 16.11] | 31.19 [28.12 – 33.80] |
| Larval development rate | 3.37 [0.88 – 8.50] | 26.37 [24.74 – 27.96] | 32.49 [30.12 – 34.81] |
| Pupal survival | 6.54 [4.00 – 8.32] | 11.23 [7.83 – 20.67] | 32.24 [28.69 – 37.38] |
| Pupal development rate | 6.05 [2.97 – 10.23] | 27.55 [25.28 – 30.59] | 33.56 [29.98 – 37.84] |
| Adult lifespan | 3.51 [2.08 – 9.09] | 17.54 [15.00 – 20.51] | 31.57 [27.36 – 35.03] |
| Fecundity | 2.72 [1.46 – 6.87] | 18.51 [17.45 – 21.51] | 34.29 [32.41 – 36.15] |
| Fitness | 7.88 [2.54 – 13.33] | 24.56 [23.07 – 25.68] | 29.41 [28.51 – 30.26] |

**Supplemental Table S6. Correlations reflecting thermal adaptation after removing the ‘POW’ population.** For correlations that were significant when including all populations (see Figure 5 in the main text), the table below shows the corresponding correlations without the ‘POW’ population. Statistically significant Pearson’s correlations (r; p < 0.05) are denoted with (*).

|  | Annual mean temp. | Warm-season maximum | # days > 35°C |
| --- | --- | --- | --- |
| $\mathrm{CT}_{\max}$ of pupal development rate | 0.535 | 0.641 | 0.691* |
| $T_{\mathrm{opt}}$ of pupal development rate | 0.565 | 0.676* | 0.705* |

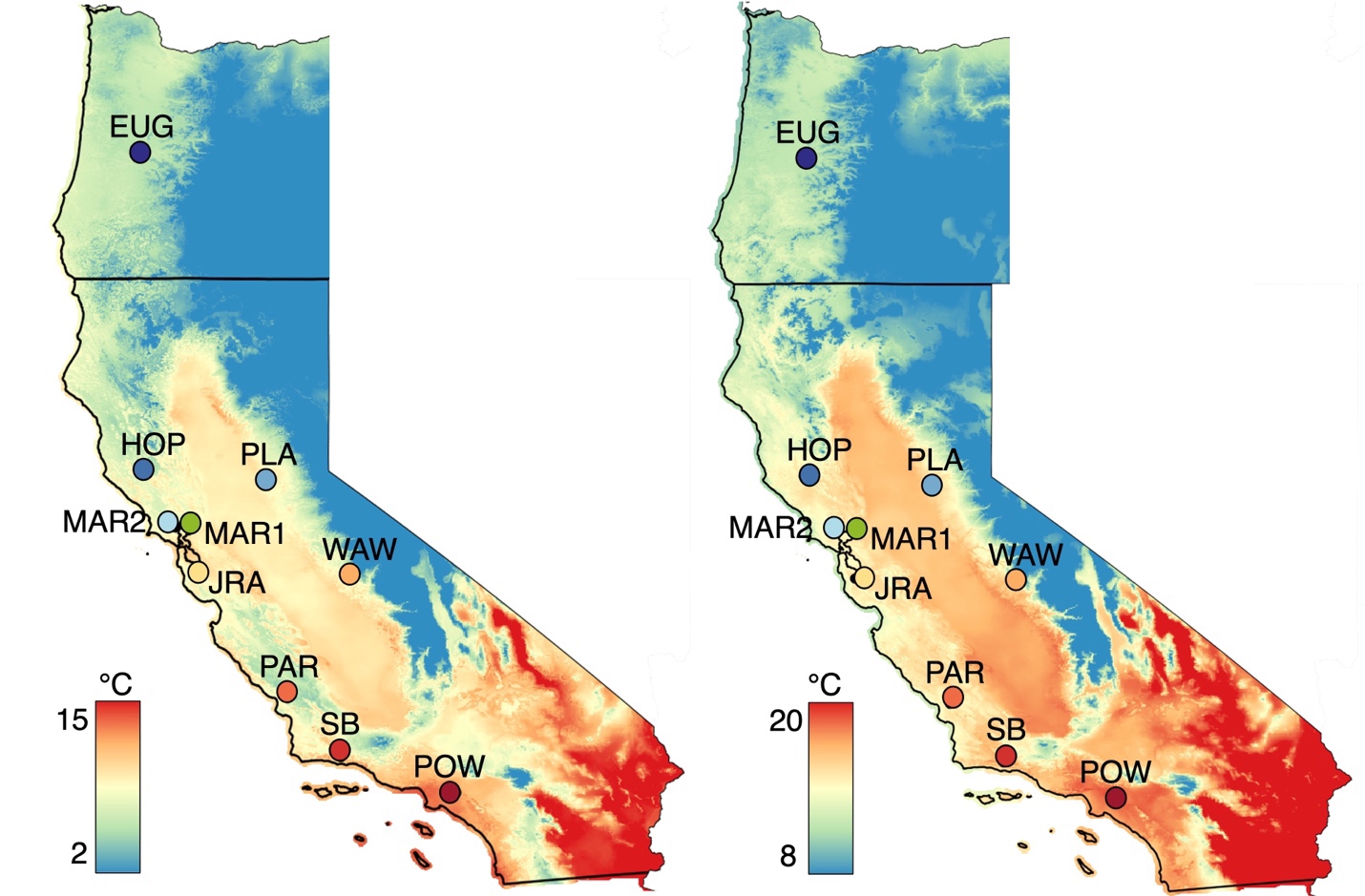

**Supplemental Figure S1.** Sample collection locations for the ten populations used in the experiment. Map colors denote the average minimum (left) and mean (right) annual temperatures (℃) from 1991 – 2020 from PRISM data. Figure 1 in the main text depicts the average maximum temperatures for this same extent and data source.

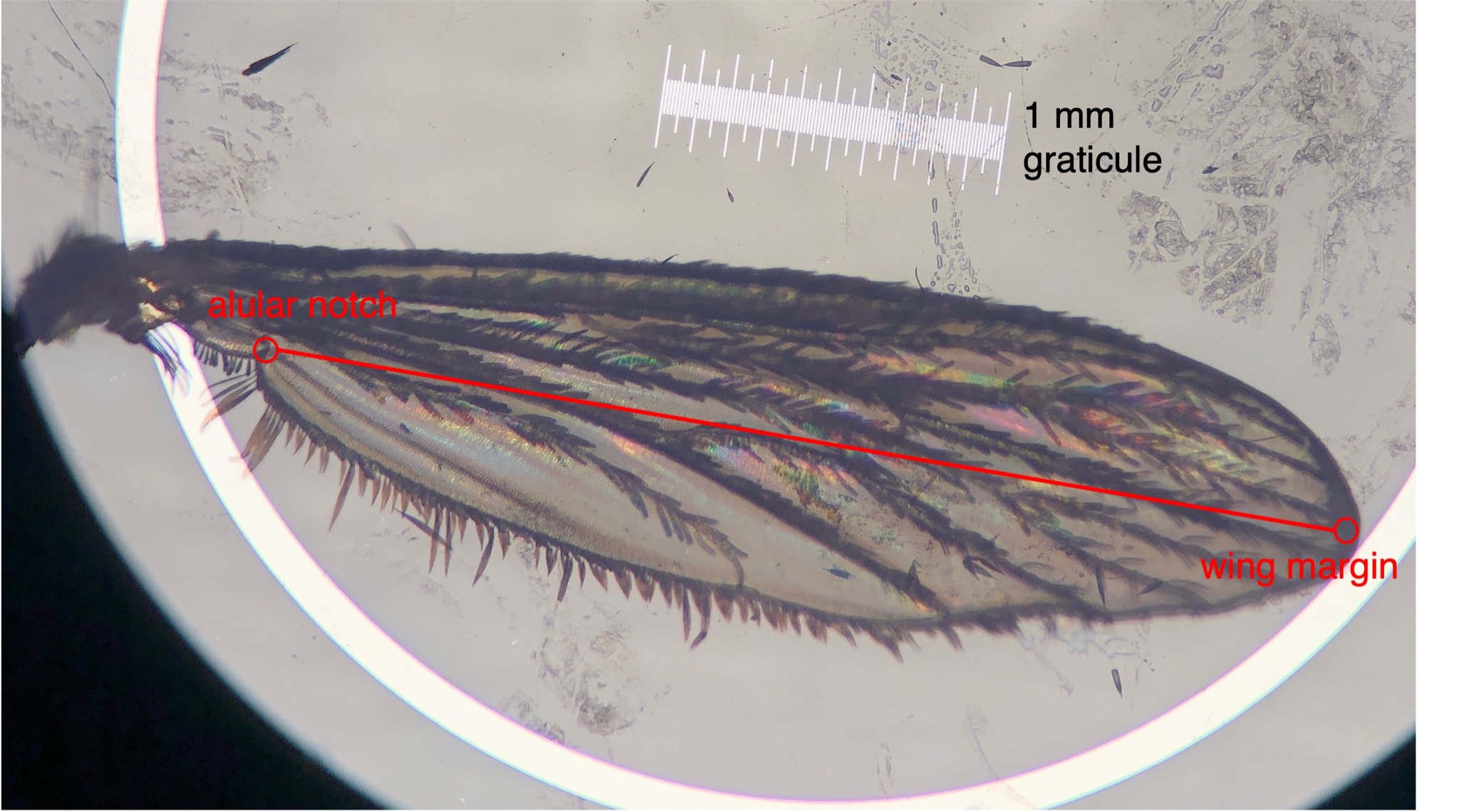

**Supplemental Figure S2.** Example wing length measurement for fecundity estimation. Wing lengths were measured as the distance from the alular notch to the tip of the wing margin, excluding the fringe scales. Measurements were made in ImageJ using a 1 mm stage graticule.

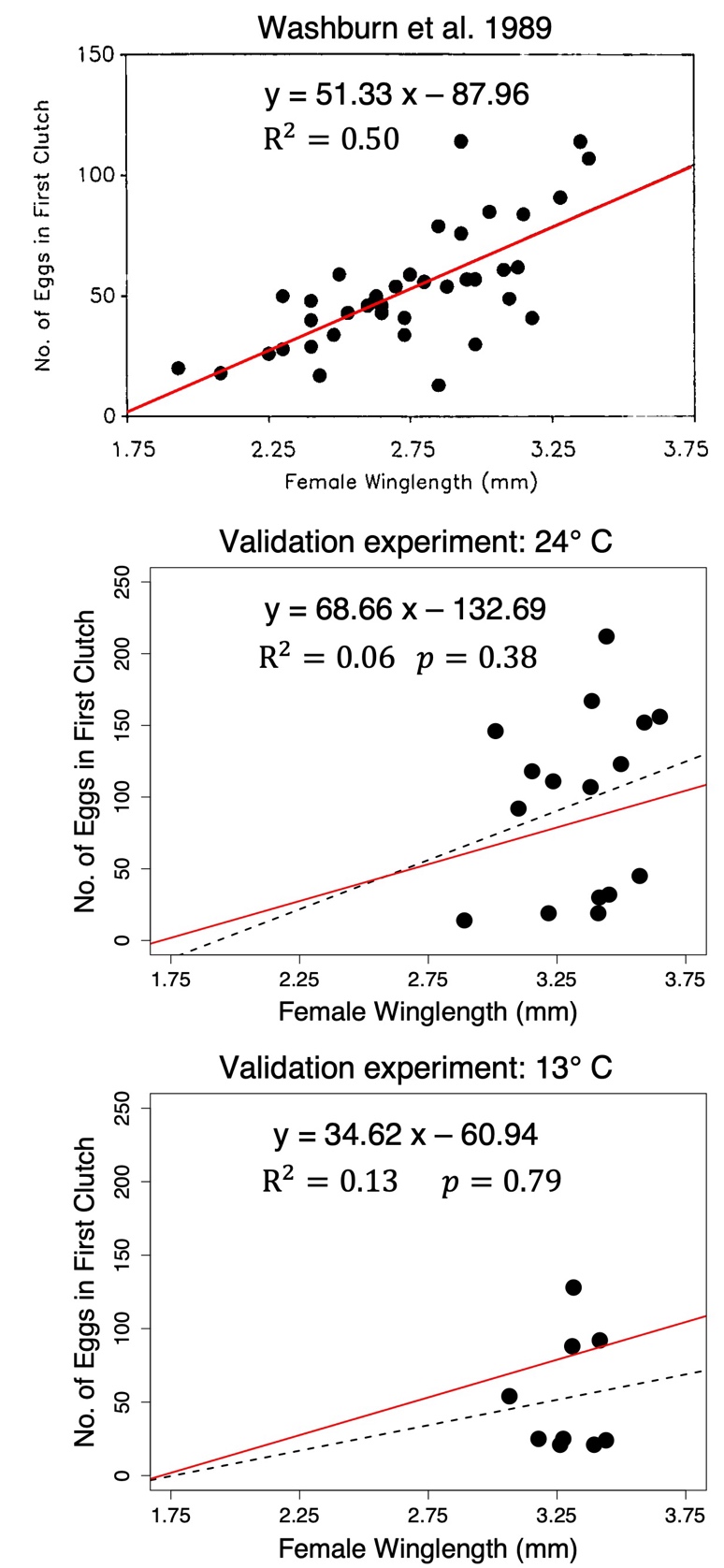

**Supplemental Figure S3.** Relationship between the number of eggs laid in the first clutch and female wing length (mm) from Washburn et al. 1989^1^ (top; reproduced from the paper), our validation experiment with blood-fed adults at 24℃ (middle) and 13℃ (bottom). Inset text at the top of each panel shows the relationship estimated from linear regression between eggs laid (y) and wing length (x; mm), and corresponding $R^{2}$ and *p* value. Solid red lines and dashed black lines denote the best fit line from Washburn et al. and our validation experiment, respectively.

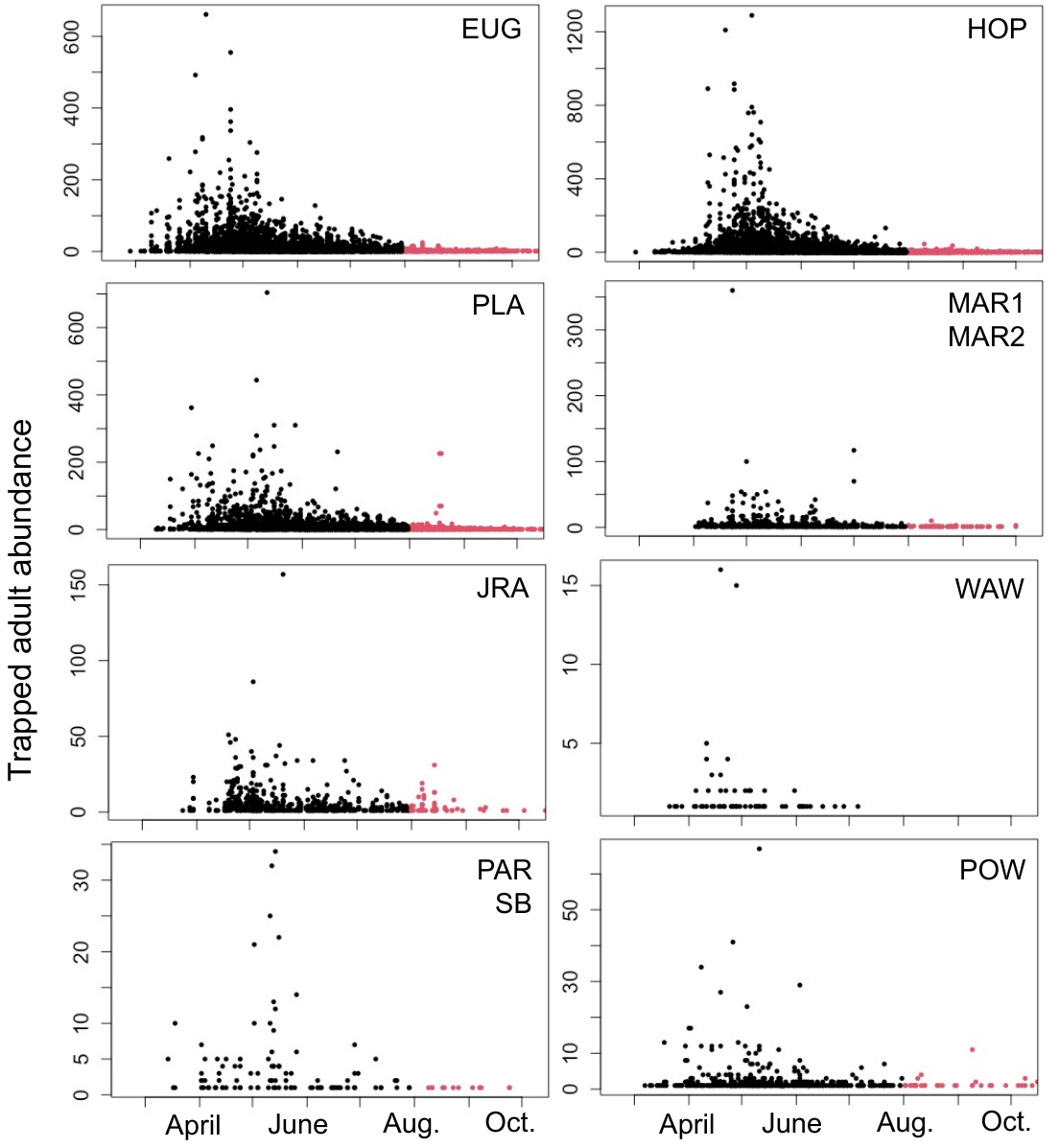

**Supplemental Figure S4.** Trap counts of adult *Ae. sierrensis* from vector surveillance near our collection sites. Data were obtained from VectorSurv and were subset to include adult trap surveillance from 2000 – 2020. Observations from before and after July 31st are colored in black and pink, respectively. The closest surveillance location for each of our field-collected populations were in: Shasta county (EUG), Lake county (HOP), Placer county (PLA), Marin county (MAR1, MAR2), San Mateo county (JRA), Fresno county (WAW), Santa Barbara county (PAR, SB), and Los Angeles county (POW).

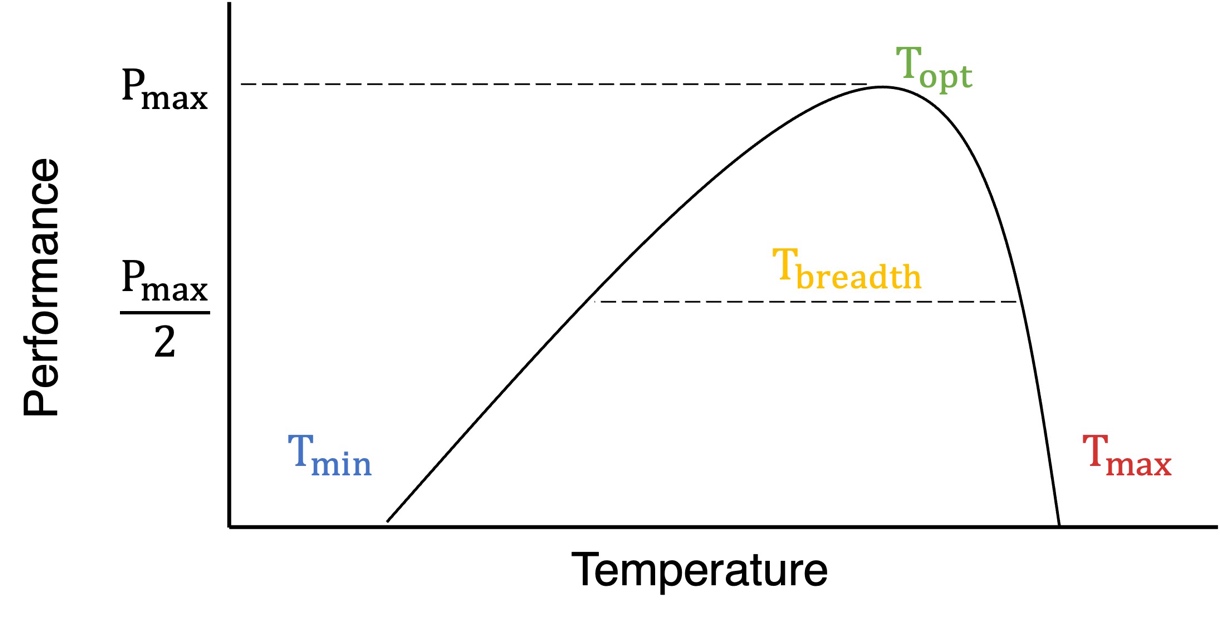

**Supplemental Figure S5.** Theoretical thermal performance curve denoting maximum performance ($P_{\max}$), the thermal optimum ($T_{\mathrm{opt}}$), the lower and upper thermal limits ($T_{\min}$ and $T_{\max}$), and thermal breadth ($T_{\mathrm{breadth}})$. T_min_ and T_max_ are the temperature values at which performance goes to zero and Topt is the temperature at which the performance peak (P_max_) is reached. Thermal breadth is calculated as the width of the curve at half of the maximum performance value (P_max_/2).

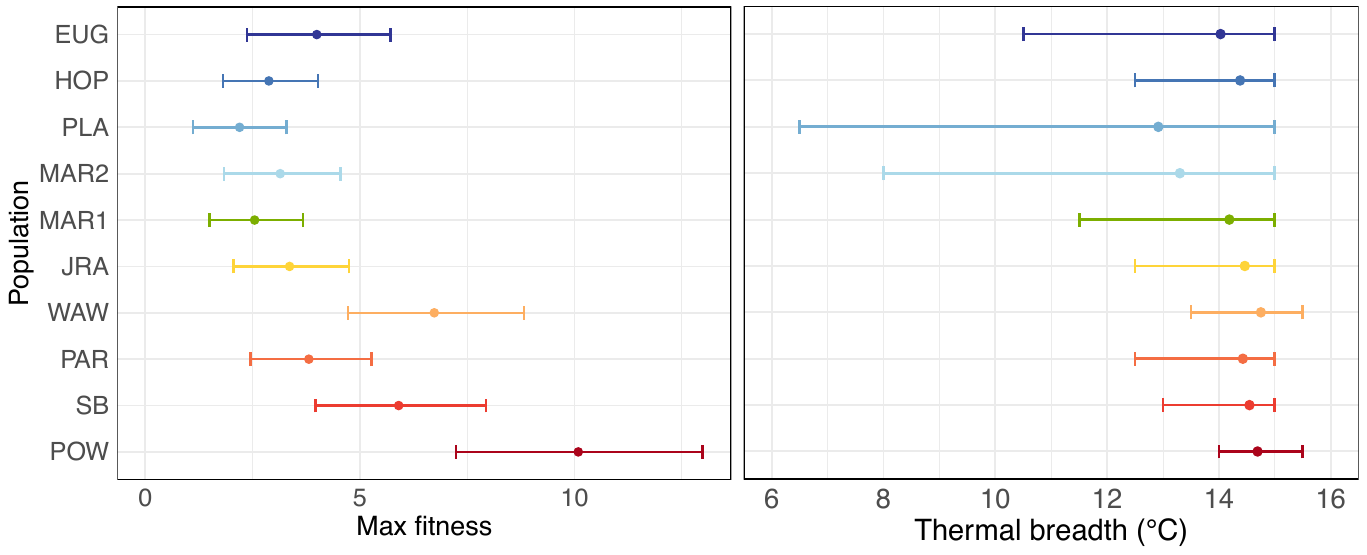

**Supplemental Figure S6. Populations vary in maximum fitness (left) but not in their thermal breadth (right).** Points denote estimated fitness thermal performance parameters for each population, and error bars denote the 95% credible intervals for each parameter. Populations (listed on the left) are colored and ordered by their latitude of collection.

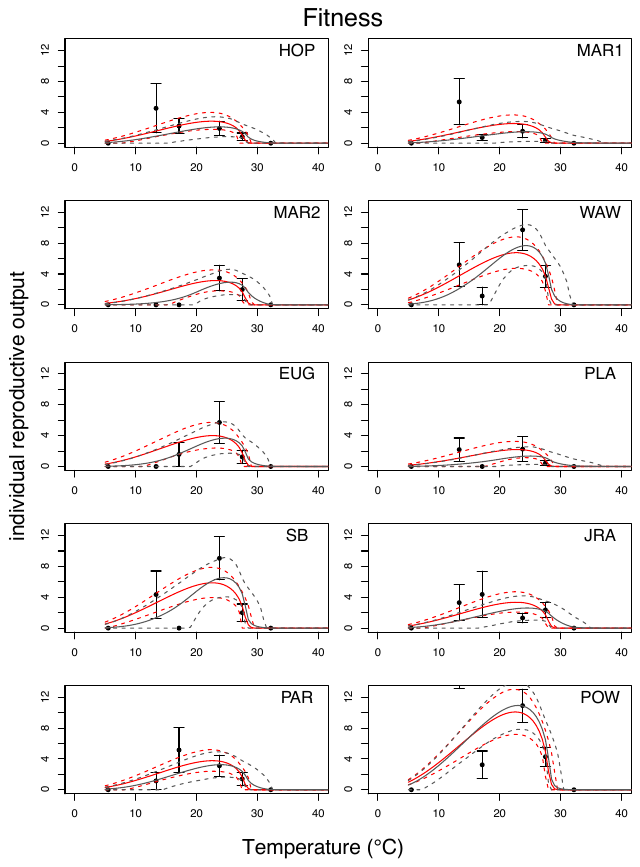

**Supplemental Figure S7.** Thermal response curves for fitness for each population. Points and error bars denote the mean and standard error from the observed data. Solid black and red lines are the mean fits from models generated using low information and informative priors, respectively. Dashed lines are the 95% credible intervals from the corresponding models. Populations, denoted in the upper right corner of each panel, correspond to the abbreviations listed in Supplemental Table S3. Note that for POW at 13°C, the mean and standard error for fitness are not visible in the plot, but are 19.14 ± 6.01°C.

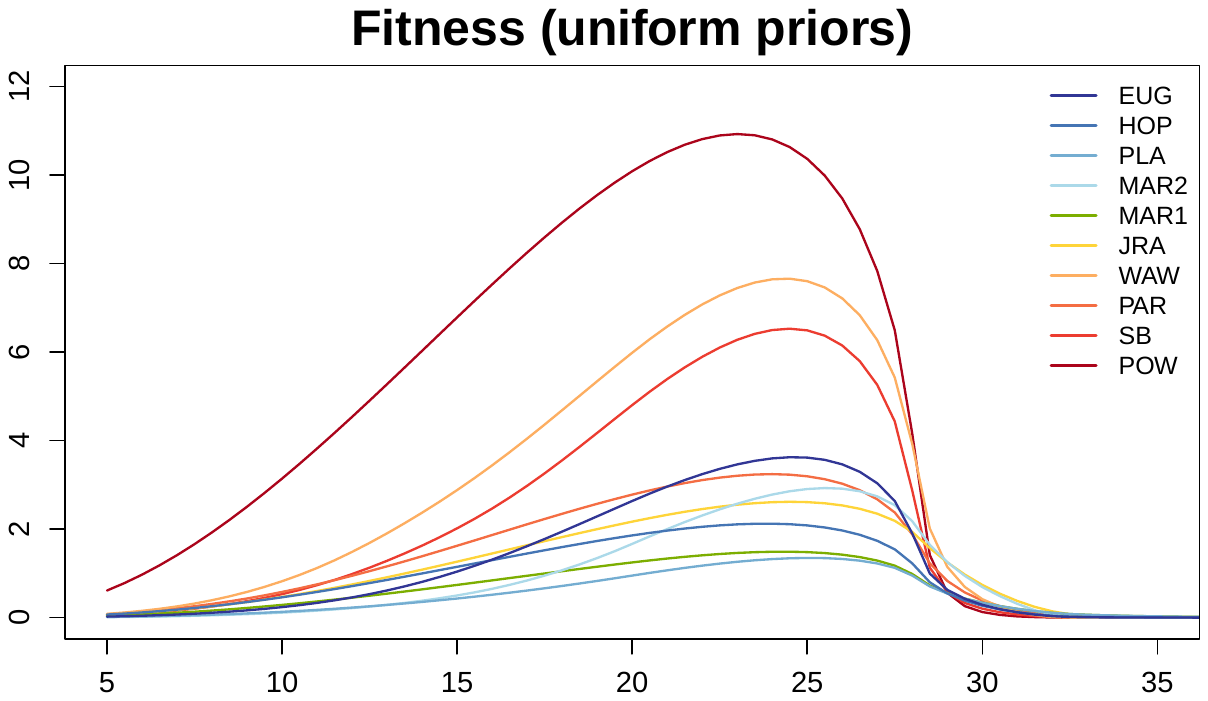

**Supplemental Figure S8.** Fitness thermal performance curves for each population fit using ‘low information’ priors. See Figure 2 in the main text for corresponding curves fit with informative priors.

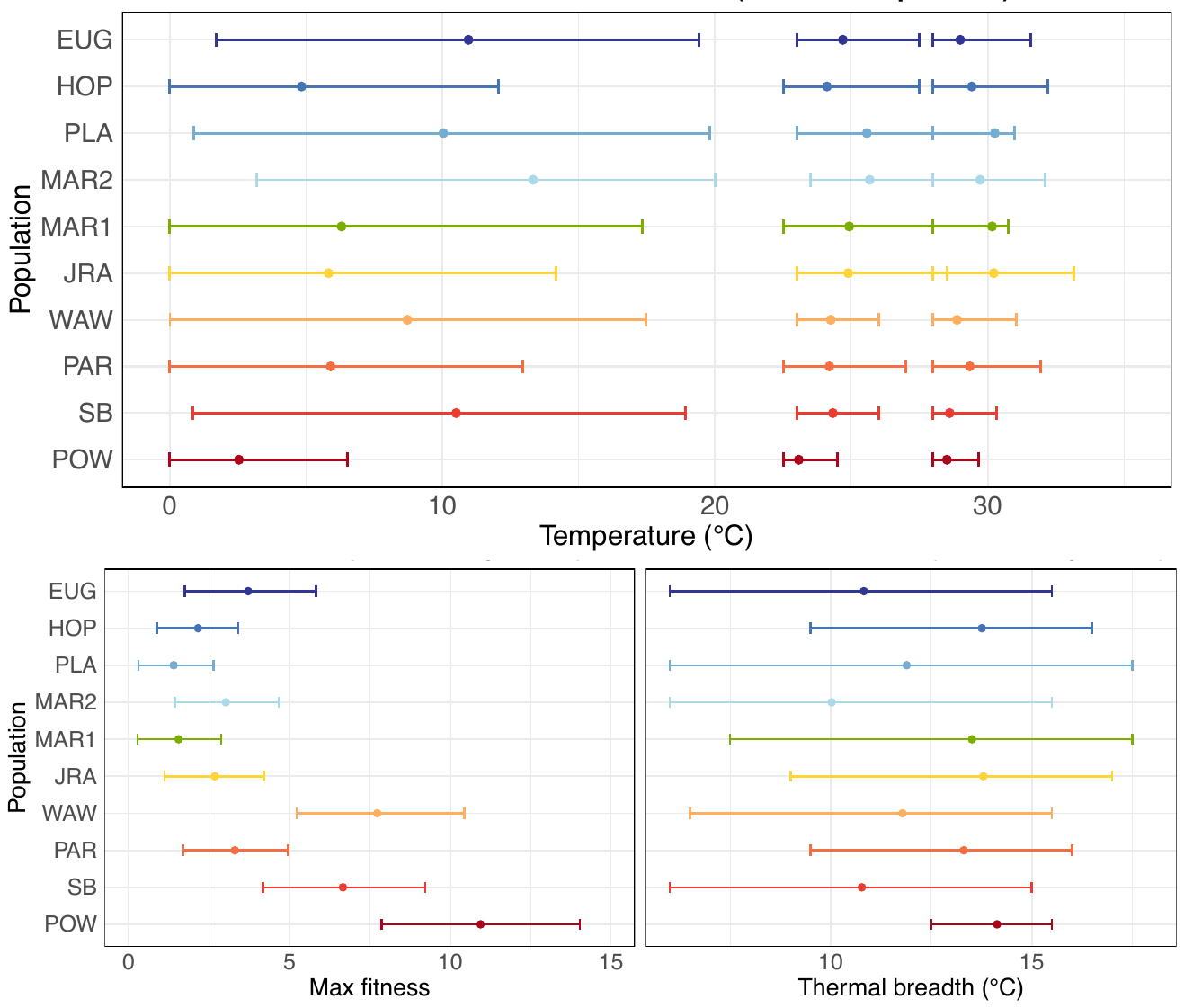

**Supplemental Figure S9.** Fitness thermal performance parameters for each population estimated using ‘low information’ priors. See Figure 3 in the main text for corresponding parameters fit with informative priors. Parameters include thermal minima, thermal optima, and thermal maxima (top), thermal breadth (bottom right), and maximum fitness (bottom left). Points and error bars denote the mean and 95% credible intervals for each parameter. Populations (listed on the left) are colored and ordered by latitude of collection.

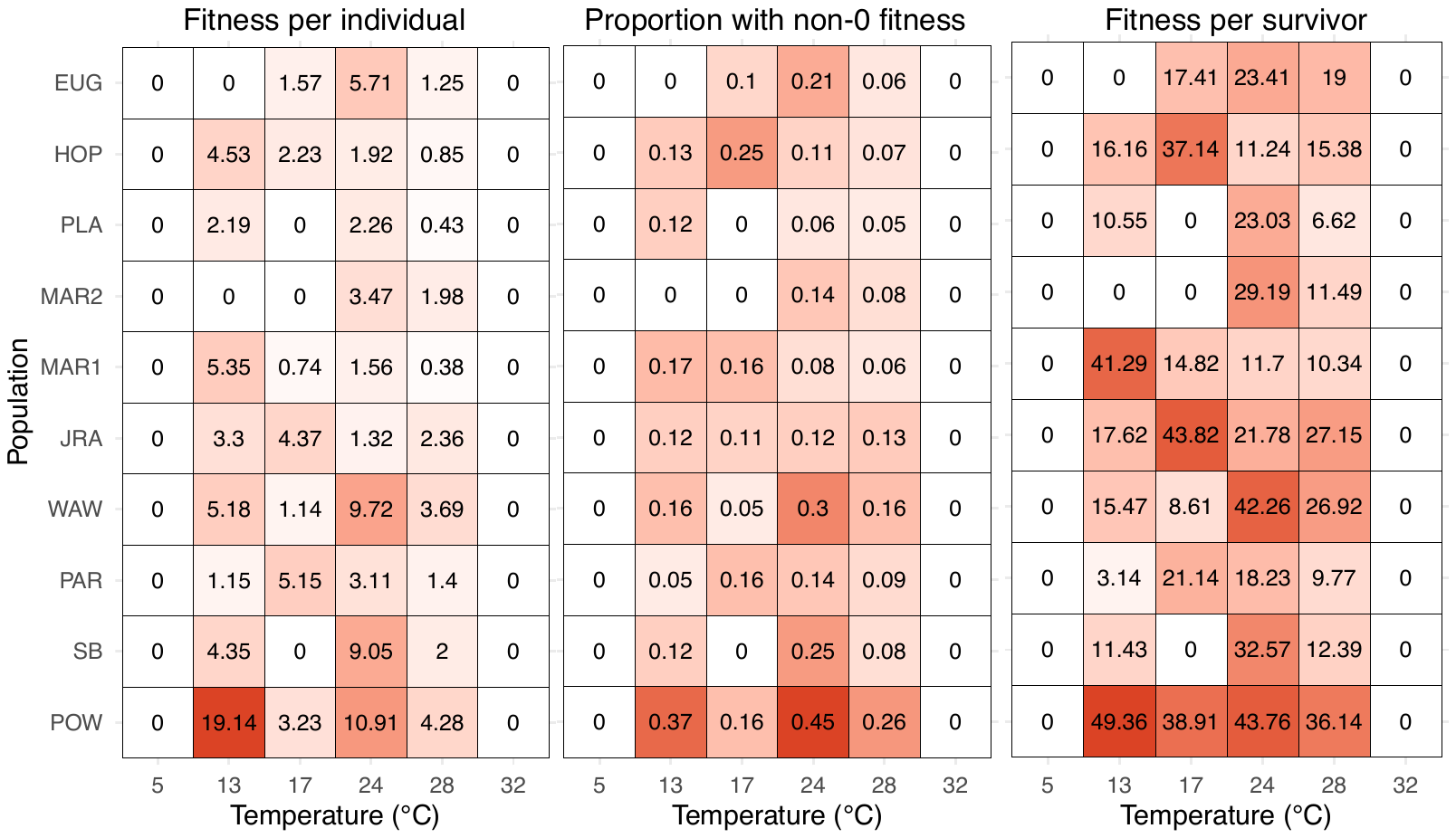

**Supplemental Figure S10.** Average fitness per individual (left), proportion of individuals with non-zero fitness (center), and average fitness per surviving individual (right) for each population and temperature treatment (bottom; units are ℃). Colors and inset values represent the same information (*i.e*., fitness values measured in different ways). Boxes colored in white represent conditions where no individuals had non-zero fitness. Note that no individuals from any population escaped larval diapause at the 5℃ treatment and no individuals survived to the date of potential oviposition at the 32℃ treatment.

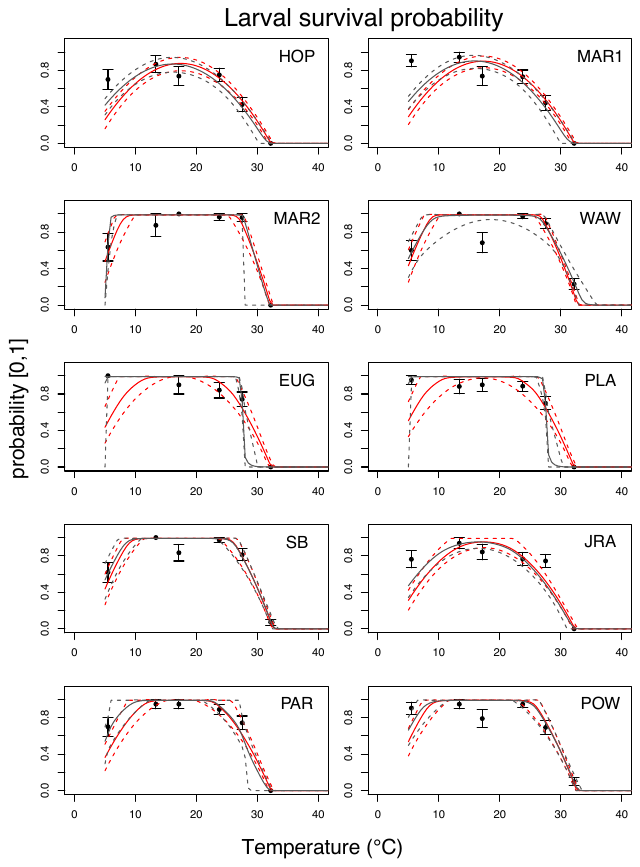

**Supplemental Figure S11.** Thermal response curves for larval survival probability for each population. Points and error bars denote the mean and standard error from the observed data. Solid black and red lines are the mean fits from models generated using low information and informative priors, respectively. Dashed lines are the 95% credible intervals from the corresponding models.

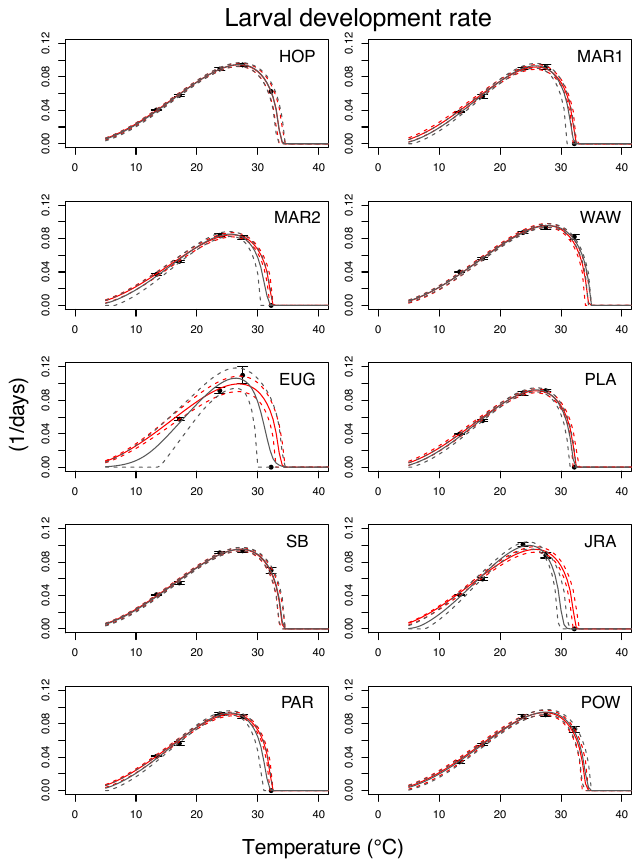

**Supplemental Figure S12.** Thermal response curves for larval development rates for each population. Points and error bars denote the mean and standard error from the observed data. Solid black and red lines are the mean fits from models generated using low information and informative priors, respectively. Dashed lines are the 95% credible intervals from the corresponding models.

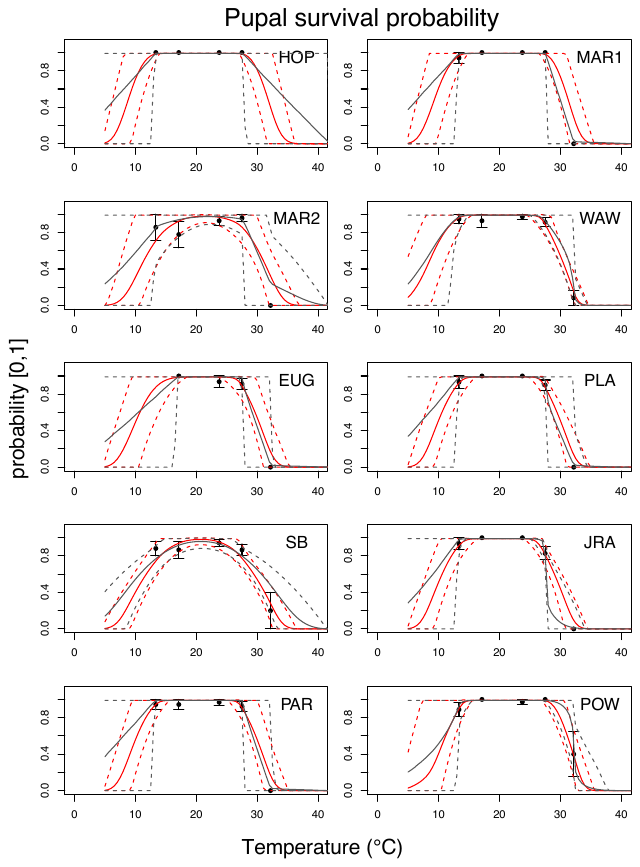

**Supplemental Figure S13.** Thermal response curves for pupal survival probability for each population. Points and error bars denote the mean and standard error from the observed data. Solid black and red lines are the mean fits from models generated using low information and informative priors, respectively. Dashed lines are the 95% credible intervals from the corresponding models.

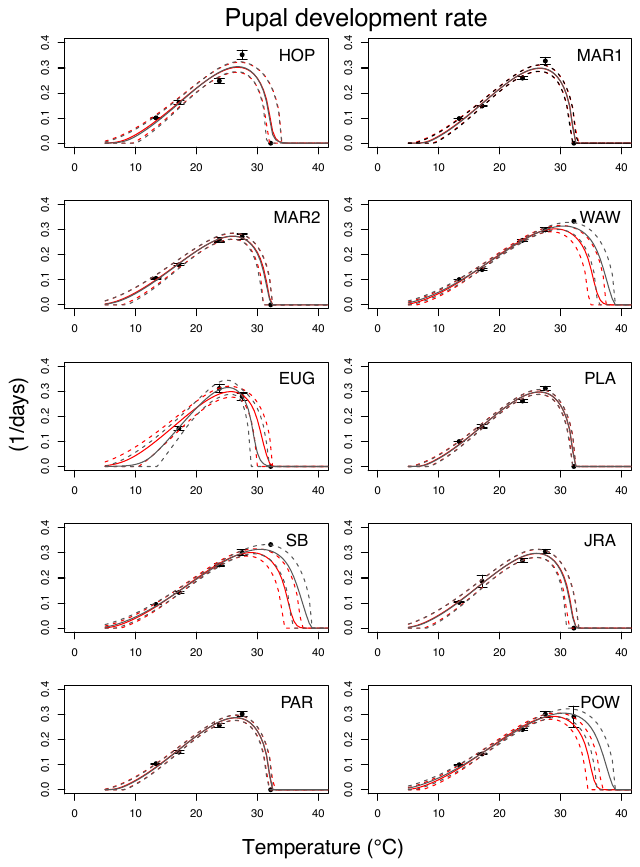

**Supplemental Figure S14.** Thermal response curves for pupal development rates for each population. Points and error bars denote the mean and standard error from the observed data. Solid black and red lines are the mean fits from models generated using low information and informative priors, respectively. Dashed lines are the 95% credible intervals from the corresponding models.

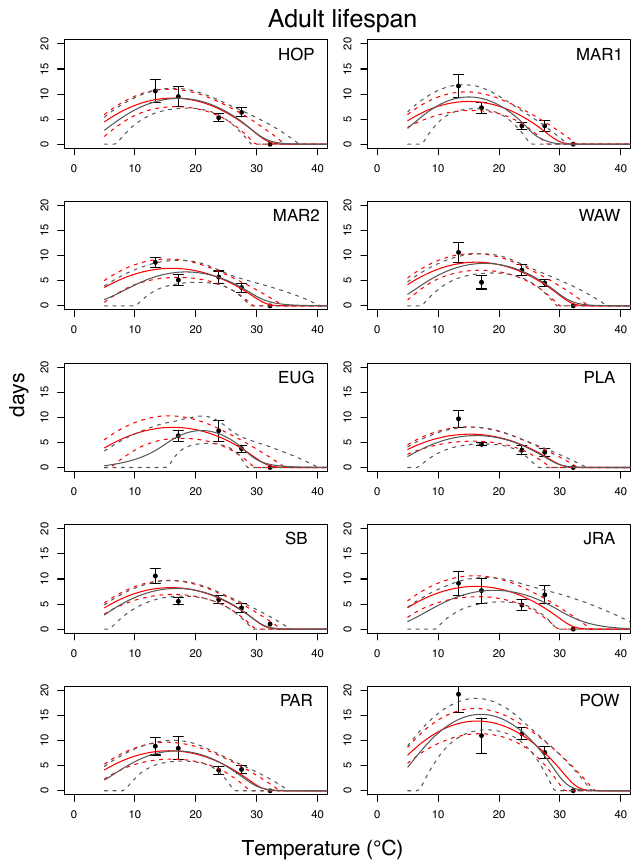

**Supplemental Figure S15.** Thermal response curves for adult lifespan for each population. Points and error bars denote the mean and standard error from the observed data. Solid black and red lines are the mean fits from models generated using low information and informative priors, respectively. Dashed lines are the 95% credible intervals from the corresponding models.

Wing length (proxy for fecundity)

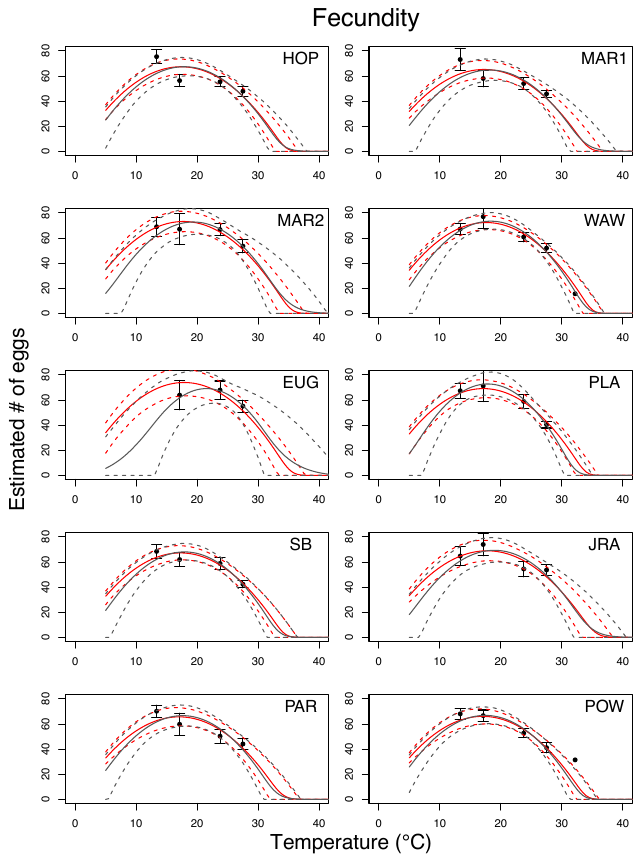

**Supplemental Figure S16.** Thermal response curves for wing length (our proxy for fecundity) for each population. Points and error bars denote the mean and standard error from the observed data. Solid black and red lines are the mean fits from models generated using low information and informative priors, respectively. Dashed lines are the 95% credible intervals from the corresponding models.

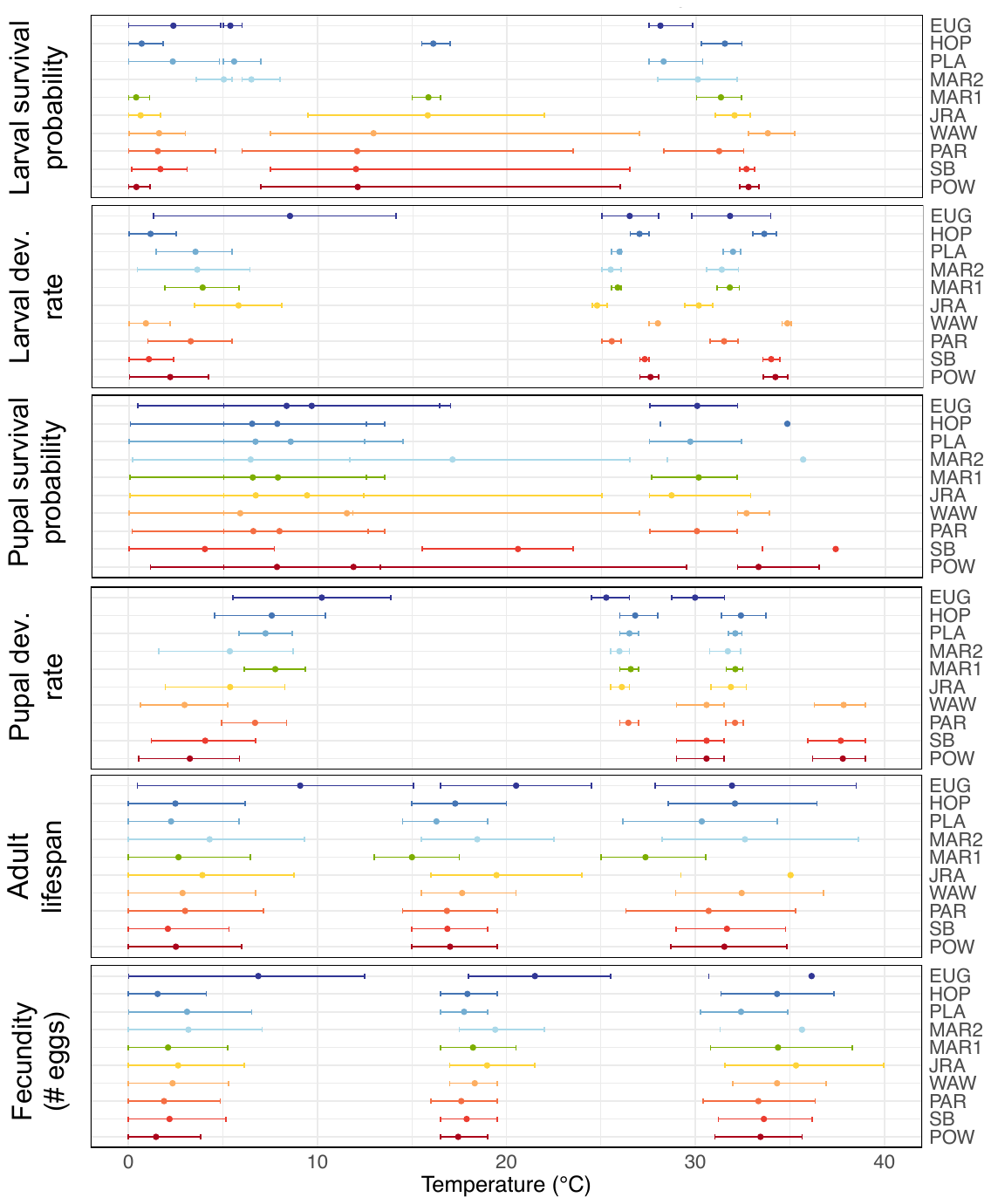

**Supplemental Figure S17.** Thermal minima, thermal optima, and thermal maxima for each life history trait and population using ‘low information’ priors. See Figure 3 in the main text for corresponding parameters fit with informative priors. Points and error bars denote the mean and 95% credible intervals for each parameter, respectively. Populations (listed on the right) are colored and ordered by latitude of collection. Units of development rates and lifespan are 1/days and days, respectively.

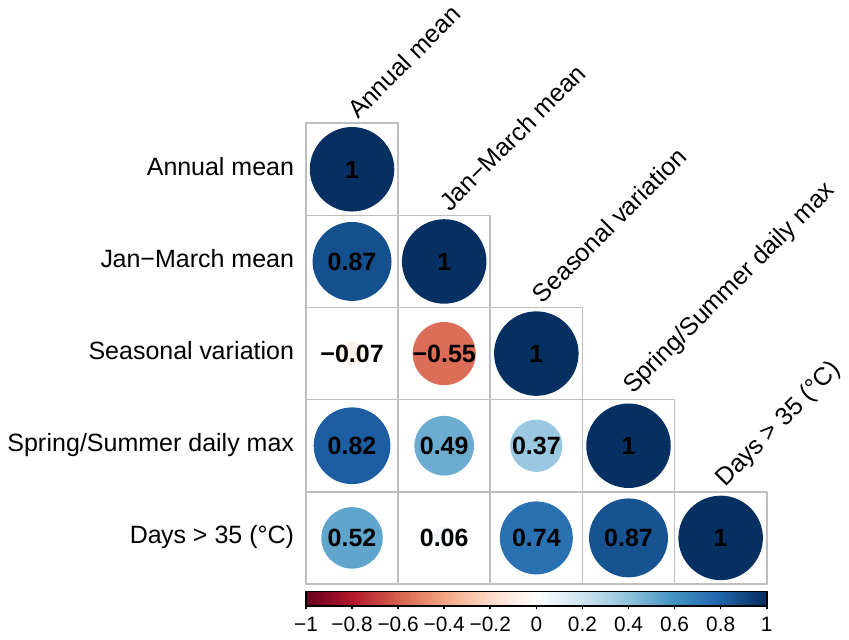

**Supplemental Figure S18.** Correlations between temperature variables characterizing the source thermal environment of each field-collected population. The size and color of the circles correspond to the strength of the correlation. For a description of how each temperature variable was calculated, see *Methods: Characterizing the source thermal environment*.

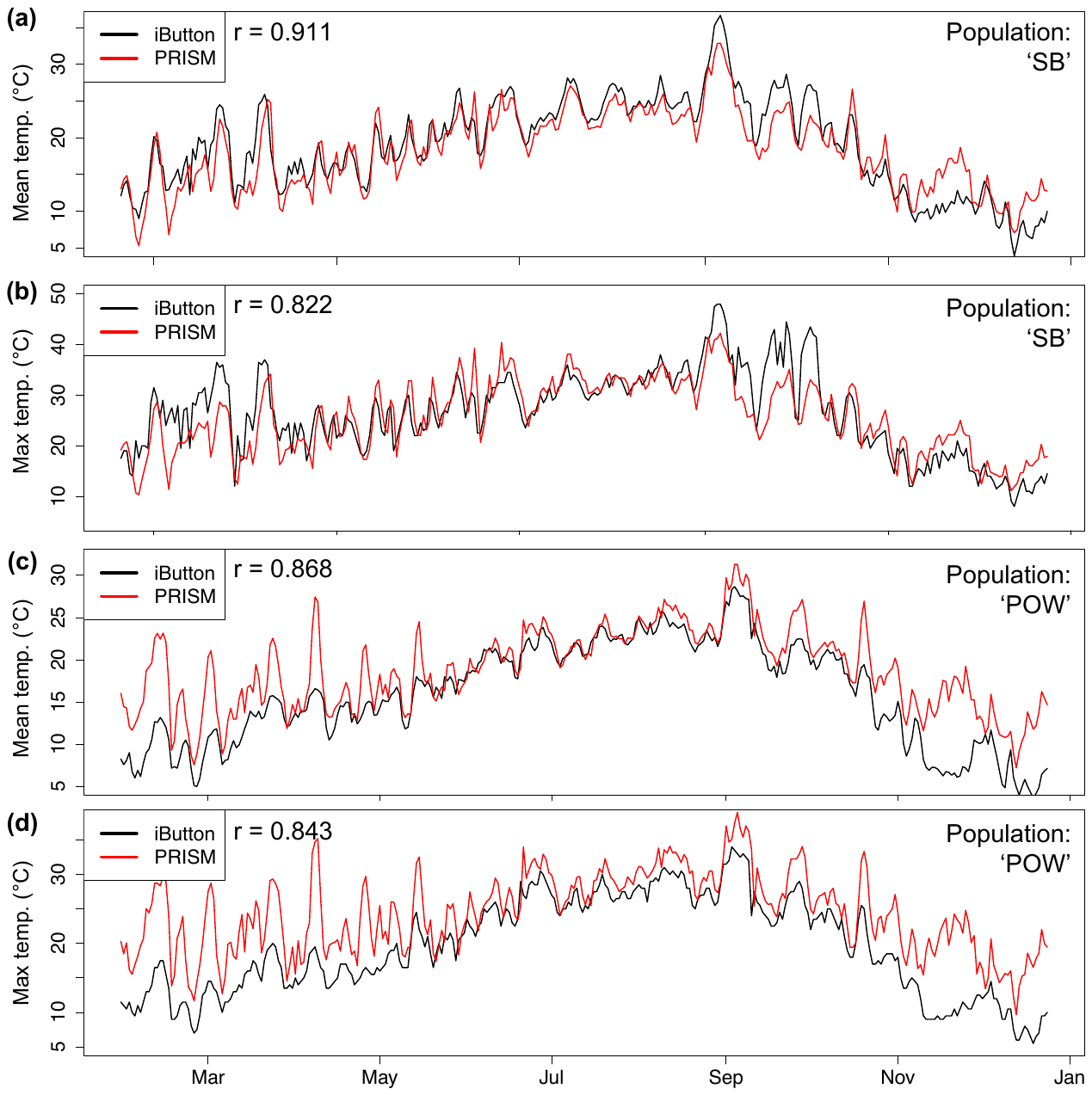

**Supplemental Figure S19.** Source temperatures measured using iButton temperature loggers (black lines) versus PRISM climate data (red lines). Values represent daily mean **(a, c)** and maximum **(b, d)** temperatures in 2022 for two populations—SB **(a, b)**, and POW **(c, d)**—the only populations for which iButtons were recovered from tree holes the year after collection. Pearson’s correlations (r) for each population and temperature metric are shown in the upper left corner of each plot.
